# The impact of cooking and burial on proteins: a characterisation of experimental foodcrusts and ceramics

**DOI:** 10.1101/2024.04.03.587902

**Authors:** Miranda Evans, Richard Hagan, Oliver Boyd, Manon Bondetti, Oliver Craig, Matthew Collins, Jessica Hendy

**Affiliations:** Department of Archaeology, University of Cambridge; BioArCh, Department of Archaeology, University of York; Independent; The GLOBE Institute, Faculty of Health and Medical Sciences, University of Copenhagen

**Keywords:** Palaeoproteomics, foodcrust, ceramics, experimental archaeology

## Abstract

Foodcrusts have received relatively little attention in the burgeoning field of proteomic analysis of ancient cuisine. We remain ignorant of how cooking and burial impacts protein survival, and crucially, the extent to which the extractome reflects the composition of input ingredients. Therefore, through experimental analogues we explore the extent of protein survival in unburied and buried foodcrusts and ceramics using ‘typical’ Mesolithic ingredients (red deer, Atlantic salmon and sweet chestnut). We then explore a number of physiochemical properties theorised to aid protein preservation. The results reveal that proteins were much more likely to be detected in foodcrusts than ceramics using the methodology employed, input ingredient strongly influences protein preservation, and that degradation is not universal nor linear between proteins, indicating that multiple protein physiochemical properties are at play. While certain properties such as hydrophobicity apparently aid protein preservation, none single-handedly explain why particular proteins/peptides survive in buried foodcrusts: this complex interplay requires further investigation. The findings demonstrate that proteins indicative of the input ingredient can be identifiable in foodcrust, but that the full proteome is unlikely to preserve. While this shows promise for the survival of proteins in archaeological foodcrust, further research is needed to accurately interpret foodcrust extractomes.

## Introduction

Proteomics has become a valuable tool for identifying food in archaeological samples particularly from exceptionally well-preserved remains from frozen [1,2], desiccated [3–5] or waterlogged contexts [6,7], as well as calcified residues such as limescale on ceramics [8,9] and dental calculus [10–13]. Due to the tissue and taxonomically specific sequence information proteins can hold, proteomics is particularly useful for the detection of ingredients and cuisine. However exceptionally well-preserved remains are rare, rendering studies with statistically significant sample sizes and regional comparisons difficult. While protein analysis of human dental calculus can be immensely valuable for understanding consumed diets, it does not necessarily give clear insights into food preparation or links with culinary material culture.

Ceramics associated with food preparation and consumption would be an ideal target sample for proteomics as they are ubiquitous in many contexts. However the detection of proteins from ceramics themselves has proved challenging, either due to strong binding, or degradation during burial. Proteins have been found to bind strongly to the mineral matrix of ceramic vessels, which likely results in good protein preservation yet renders their extraction challenging [14] without the use of harsh solvents [15]. Conversely, Barker et al. [16] concluded that protein content decayed rapidly upon burial, although they did not measure the initial protein content prior to burying their samples, and thus the rapidity of protein loss may be difficult to estimate. Food proteins have been reported from archaeological ceramics [8,17–19] and modern replicas [20,21]. However there are potential factors aiding the detection of proteins in archaeological cases, such as the inclusion of remnant encrustations [17], or the sampling of ceramic from immediately beneath a limescale deposit [8], which may have provided protection from diagenesis. In the case of Solazzo et al. [18], the sherd was from relatively cold conditions in the Arctic coast of Alaska, and contained lipid-rich foods including whale and seal meat, both factors which may have improved protein preservation, although we note that food proteins were not detected in similar ceramics in a later study [22].

### Biomolecular analyses of foodcrusts

Given the challenges in extracting proteins from ceramics themselves, foodcrusts may offer a good alternative target sample for proteomic analysis. Foodcrust, sometimes referred to as “carbonised residue” and “char” is broadly defined as “amorphous charred or burnt deposits adhering to the surface of containers associated with heating organic matter” [23] The prevalence of foodcrusts varies considerably, however they are particularly abundant in Mesolithic and Neolithic contexts in Northern Europe and Eurasia where they are sometimes found on the majority of ceramics within assemblages [24,25].

Lipid analysis of foodcrusts has considerably improved our understanding of ancient diet and particularly of marine resource utilisation. The method has been applied to detect food in assemblages across a vast geography spanning Europe and Northern Asia [24,26–32] East Asia [25,33–37] and the Americas [22,38], and to select samples that do not contain aquatic resources for use in carbon dating, which are thus unhindered by the reservoir effect [23]. The formation of foodcrusts is a topic of ongoing research. They are often presumed to be formed by cooking food, although they can also result from the use of fuel for illumination [39] or the production of sealants, moisturisers, adhesives or glues [23,40,41]. A possible correlation exists between foodcrust formation and the processing of aquatic resources, or alternatively these particular lipids may preserve better in foodcrusts than ceramics [24].

Proteomic analysis has recently been applied to archaeological foodcrusts, demonstrating the viability of the technique [42,43], but also has generated questions around protein survival and biases [42,43]. Results so far appear congruent with the association between aquatic resource processing and foodcrusts. Shevchenko et al [43] performed proteomic analysis on four Mesolithic-Neolithic foodcrusts from the site of Friesack 4, Germany. Their results revealed the presence of deamidated fish vitellogenins and parvalbumins in one foodcrust and *Suidae* collagens detected in another. Lyu et al [42] analysed 21 foodcrusts from the site of Xiawan in South-East China for both lipids and proteins. Their results revealed the presence of potential dietary proteins in five samples, including myosin from large yellow croaker and mandarin fish, and collagen from Caprinae and potentially other mammals [42]. In both of these cases a low proportion of samples analysed produced dietary results, and the number of dietary proteins and peptides was also low. Despite this initial headway, many questions remain outstanding concerning the survival of proteins in ancient foodcrusts.

### Potential preservation biases

The key question is simply the degree to which the proteins identified in ceramics and their residues reflect the original food processed in the vessel. Although proteins indeed become altered through different cooking processes, we remain ignorant of the degree to which those changes impact the detection of proteins in ceramics and foodcrust residues. Similarly, we are also unaware of the impact of burial on the survival of food proteins in these samples.

Compared to proteomics, there is a diversity of published experiments exploring how lipids derived from different ingredients respond to a range of cooking and deposition practices [for example 44,45–50]. For instance, Miller et al. [44] demonstrate that absorbed lipids extracted from ceramics represent a long period of use, while surface deposits represent the most recent cooking events. However, as a much younger discipline, such studies are rarer in palaeoproteomics with most experimental studies focused on understanding if proteins survive at all in ceramics, or optimising extraction protocol [15,16,21,51,52] rather than the range of cooking and deposition variables which may impact them [although see 3,9].

In this study, we characterise the impact of cooking and entrapment in foodcrust and ceramic, followed by burial on the identification of proteins and peptides from three common Mesolithic foods: *Cervus elaphus* (red deer), *Salmo salar* (Atlantic salmon) and *Castanea sativa* (sweet chestnut), to anticipate results and consider expectations for archaeological interpretations of diet and food preparation practices derived from foodcrusts and ceramics. We aim to identify proteins that persist throughout the cooking process and become embedded in foodcrusts and ceramics, as well as those that persist through burial in soil for six months. Specifically, we examine metrics including peptide and protein count, concentration of different amino acids, peptide length, peptide hydropathicity, peptide isoelectric point, protein thermal stability, protein secondary structure, protein disorder, protein amyloid propensity and relative solvent accessibility of the detected peptides in order to identify characteristics of their survival, and we also compare protein with lipid content. Given that proteomics is capable of providing highly specific information concerning ancient ingredients and cuisine, understanding the impact of cooking and burial on the protein content of ceramics and foodcrusts is crucial to accurately interpreting proteomic results of ancient samples. These results provide a maximum baseline for protein recovery from ceramics and foodcrusts under similar conditions.

## Materials and methods

### Sample creation

Experimental samples included deer meat, salmon meat and chestnut flour, each individually cooked in replica ceramic vessels to induce foodcrust formation, before being split and one half buried (Figure 1, see also supplementary text S2). These were originally created and described previously by Bondetti et al [47] to investigate the formation and diagnostic value of ω-(o-alkylphenyl)alkanoic acids (APAAs). After lipid extraction, samples were stored at 4°C until protein analysis was performed in summer 2020. Full details of the methodology used for sample creation, protein extraction, and machine analysis can be found in the electronic supplementary material S1.

**Figure 1:**
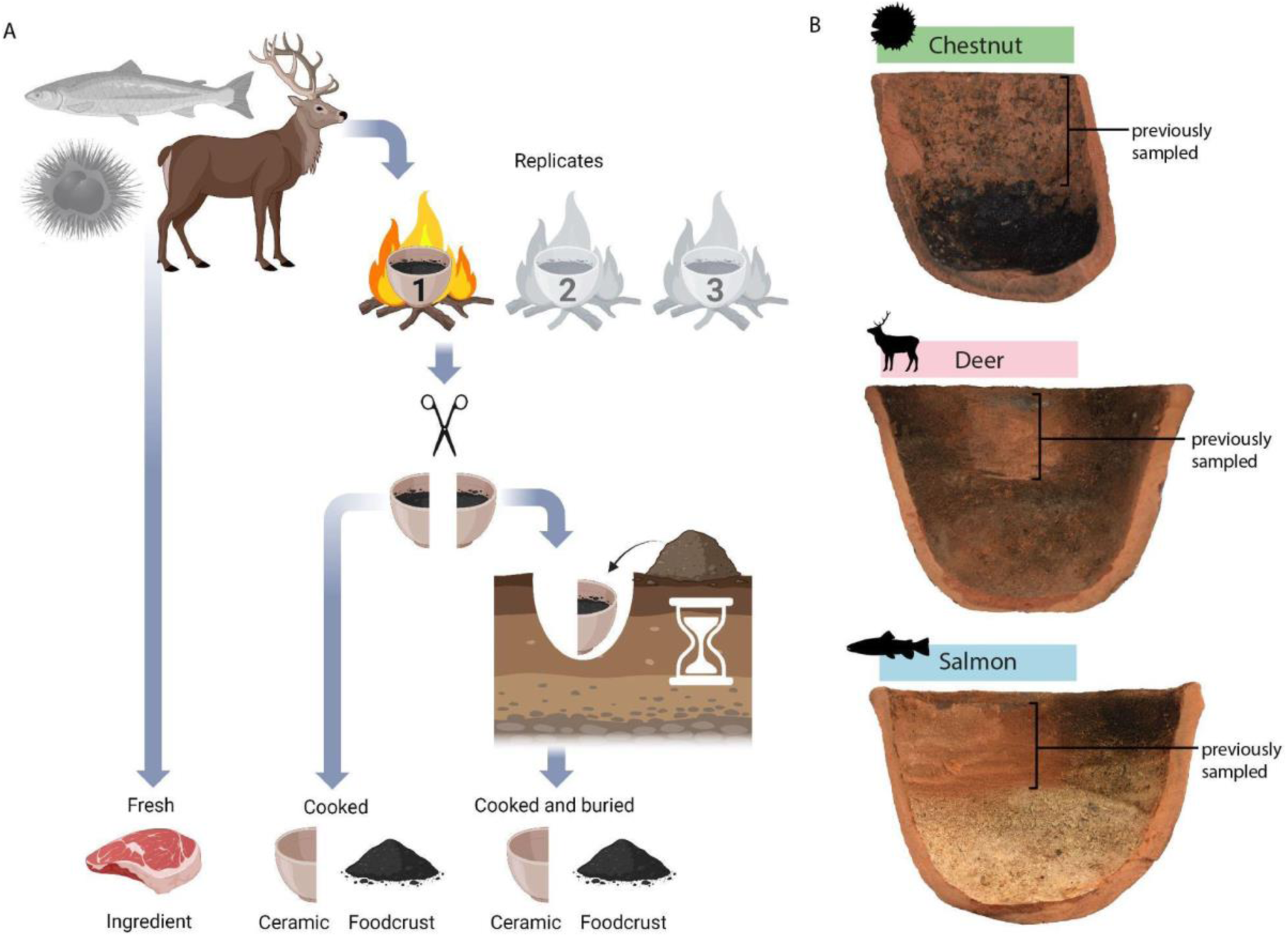
A: Sample creation process (Image created with BioRender.com), B: Example of cooked foodcrust samples. Top: Chestnut, Middle: Deer, Bottom: Salmon.

### Protein extraction

All samples and machine washes were analysed following an SP3 protein extraction protocol [53,54] adapted for ancient samples [55,56] which can be found on protocols.io [https://doi.org/10.17504/protocols.io.bfgrjjv6] and is routinely applied to archaeological samples [56–58].

### LC-MS/MS analysis

The samples were analysed on an Orbitrap Fusion at the Centre for Excellence in Mass Spectrometry at the University of York. Blank machine washes were run between each sample injection in order to examine and reduce the degree of carry-over between samples.

### Data analysis

Samples and machine washes were analysed using Maxquant (version 2.1.0.0). Peptides were searched allowing for tryptic cleavage, minimum length of seven amino acids, with both a protein and peptide target false discovery rate of 1%. Variable modifications included oxidation (M), acetylation (protein N-term), deamidation (NQ), glutamine to pyroglutamic acid, and the fixed modification of carbamidomethyl (C) was specified.

All samples were searched against a combined database which included an European Red Deer proteome (UP000242450), an Atlantic salmon proteome (UP000087266), a Chinese chestnut proteome (UP000737018) and “cRAP”, a database of common lab contaminants. *Castanea mollissima* (Chinese chestnut) was chosen as a reference database as there was no proteome for *Castanea sativa* (Sweet chestnut) at the time of analysis. To investigate any potential cross-contamination during the outdoor cooking experiment, in the lab and due to carry over in the LC-MS/MS, all samples were searched against all databases, to establish a baseline of cross-contamination. The ‘match between runs’ option was not allowed, given the varying proteomes present as match between runs has been found to falsely inflate peptide identifications [59]. Lowest common ancestor (LCA) was generated for peptides where possible using Unipept Desktop (version 2.0.0). The data was filtered to remove potential machine carry-over and laboratory contaminants. Full details of this process can be found in supplementary text S1. It became apparent that cross contamination occurred during field experiments, which is particularly evident in low protein samples such as ceramic extracts (supplementary text S7). To minimise the impact of cross contamination on protein characterisation, only samples with known cross-contaminant peptide concentration ≤2% of the total peptide count were included in the analysis of protein properties.

## Results and discussion

### Impact of cooking and burial on protein and peptide detection in ceramics and foodcrust

#### Do proteins preferentially survive cooking and burial in ceramics or foodcrust?

It is immediately apparent that foodcrusts are more likely to harbour preserved proteins than ceramics using the extraction methodology utilised here. Overall, the peptide and protein count for each food was high in the fresh ingredient, reduced in the cooked foodcrust, and further reduced (yet still appreciable) in the buried foodcrust samples, with some variation depending on ingredient (Figure 2, see also supplementary text S3) (i.e. <4-8 proteins in chestnut foodcrusts, <24-27 in salmon foodcrsuts, <16-33 in deer foodcrusts, supported by >1 peptide spectral match). In contrast to the foodcrust, protein and peptide counts were much lower in the ceramic samples, even prior to burial, and remained extremely low after burial (Figure 2). This indicates that small but appreciable numbers of food proteins may be detectable in foodcrusts in archaeological samples of similar ingredients and conditions but in contrast, given that few positive protein identifications could be made from buried ceramics, ceramic samples should not be expected to result in positive protein results if a similar protocol is followed.

**Figure 2.**
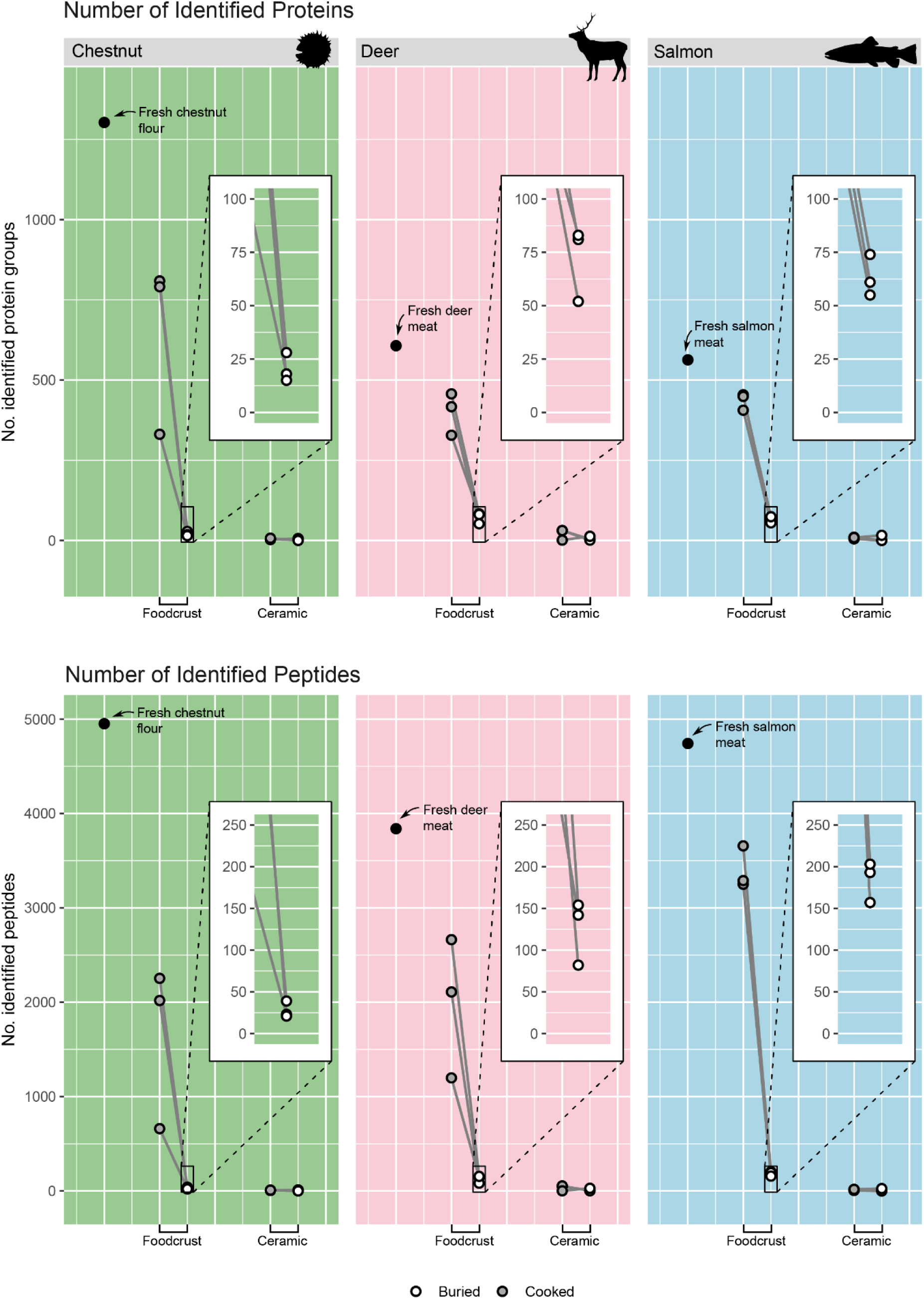
The number of identified proteins (top) and peptides (bottom) in foodcrust and ceramic samples cooked with chestnut, deer and salmon. Counts include all peptides/proteins not excluded by filtering steps described above.

#### Why are so few proteins detected in ceramics?

Euclidean clustering revealed that the cooked ceramic samples clustered together with the buried cooked ceramic samples for all food types (Figure 3A), indicating a strong correlation between the cooked ceramic sample and the buried cooked ceramic sample. This likely demonstrates that the proteins were too strongly bound to the ceramic matrix to be extracted using the protocol here employed, or alternatively, that even prior to burial, protein identification was already very low in ceramic samples, for instance if protein content had not yet impregnated the ceramic walls or if cooking had rapidly degraded any protein content present in the ceramic. This sharp reduction of both protein and peptide count in the ceramic samples compared to the fresh ingredients, is unsurprising in light of existing published research indicating similar findings [14], and that positive results from ceramics lacking encrustations and under normal preservation conditions have rarely been reported [although see 18,19,20]. Further work is necessary to devise optimal extraction methods for ceramic-bound protein.

**Figure 3:**
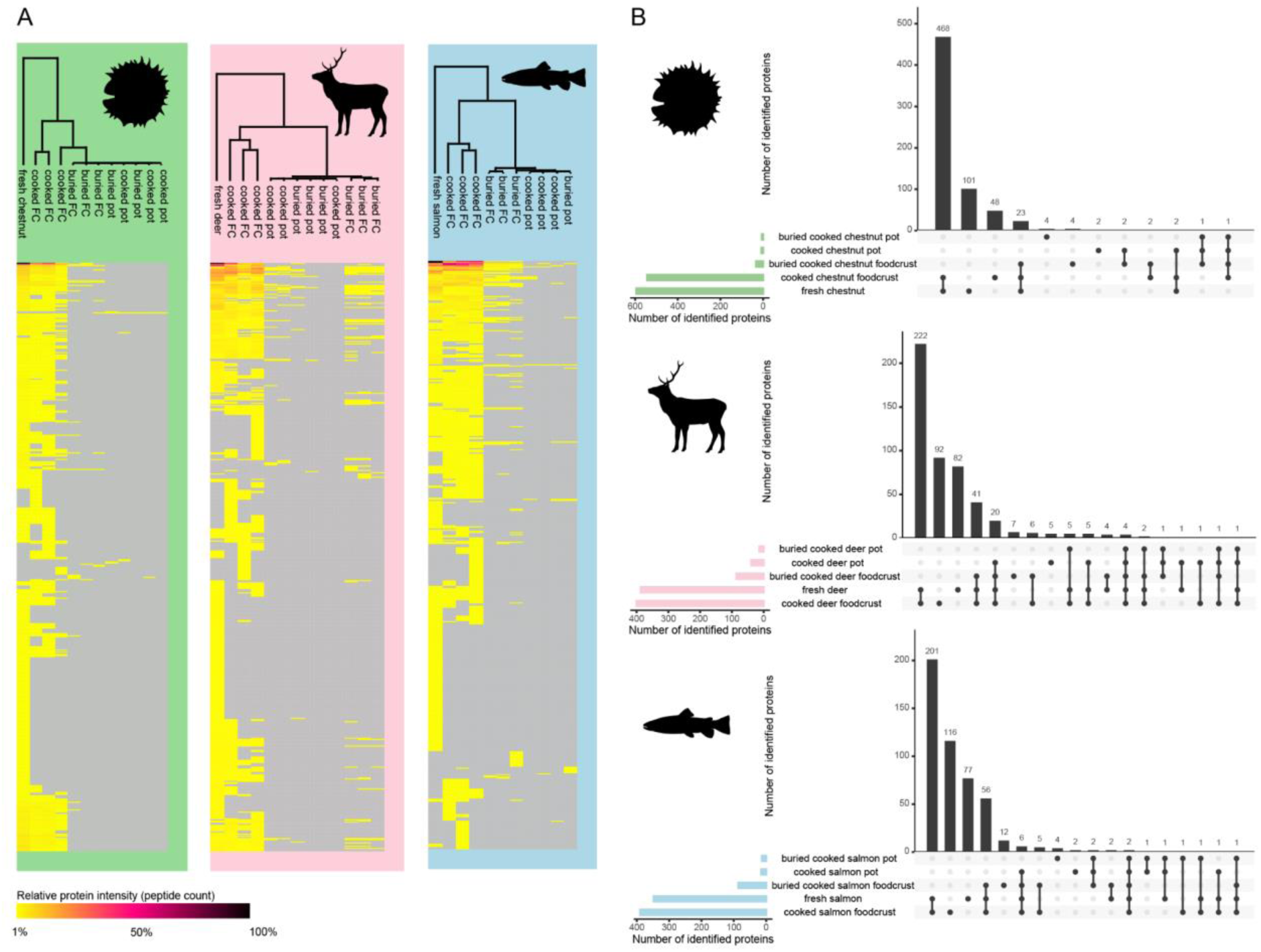
A: Hierarchical cluster analysis of proteins identified in all chestnut (left), deer (middle) and salmon (right) foodcrust and ceramic samples (Euclidean correlation), created in Perseus version 1.6.14. B: Upset plot displaying intersection of proteins observed in ceramic, foodcrust and fresh sample categories for chestnut (top), deer (middle) and salmon (bottom) samples. Created using UpsetR [60]

#### Which proteins survive cooking and burial?

A key aim of this study was to identify the proteins that persist throughout the cooking process and become embedded in foodcrusts and ceramics, as well as those that persist after burial for 6 months. The most abundant proteins detected in foodcrust and ceramics (by peptide count) can be seen in Table 1. The most abundant proteins detected in foodcrusts surviving cooking and burial for 6 months can be seen in Table 2. We note that the most abundant proteins detected in ceramic samples include several probable contaminants such as keratins, while the most abundant foodcrust proteins tend to contain more genuine ingredient matches. We also note that despite filtering described above, some cross contamination is observable especially in chestnut samples, likely derived from the field experiments.

**Table 1.**
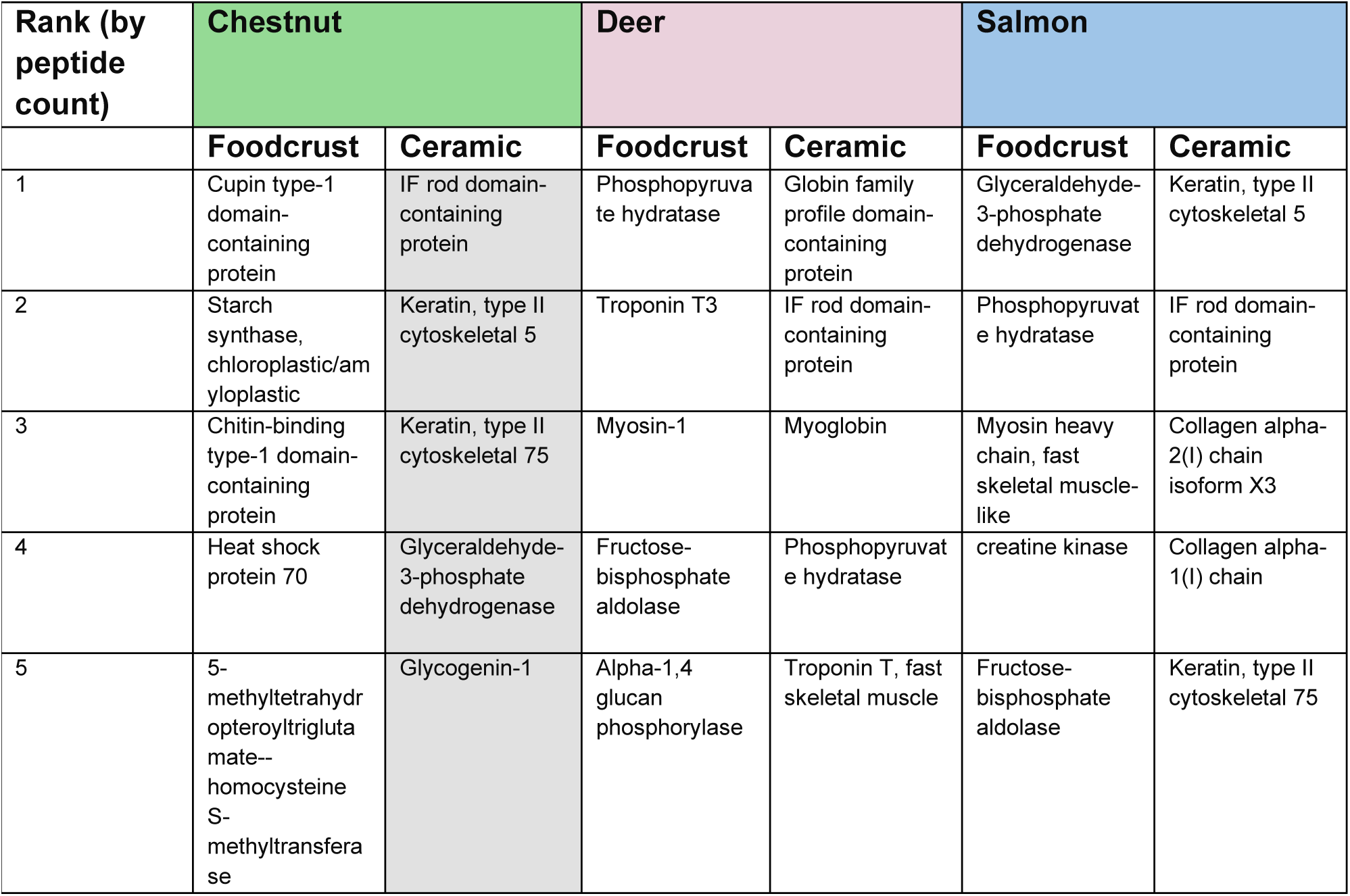
The top 5 most abundant proteins (by peptide count) preserved in foodcrust and ceramic samples after cooking. Data from replicates has been merged. Grey indicates likely contaminant or cross-contamination.

**Table 2.** The top 5 most abundant proteins (by peptide count) preserved in foodcrust and ceramic samples after cooking and burial for 6 months. Data from replicates has been merged. Grey indicates likely contaminant or cross-contamination.

| Rank (by peptide count) | Chestnut |  | Deer |  | Salmon |  |
| --- | --- | --- | --- | --- | --- | --- |
|  | Foodcrust | Ceramic | Foodcrust | Ceramic | Foodcrust | Ceramic |
| 1 | TATA box binding protein associated factor (TAF) histone-like fold domain-containing protein | IF rod domain-containing protein | ACTC protein | L-lactate dehydrogenase | Glyceraldehyde-3-phosphate dehydrogenase | IF rod domain-containing protein |
| 2 | IF rod domain-containing protein | Keratin, type II cytoskeletal 5 | Globin family profile domain-containing protein | Collagen alpha-2(I) chain | Myosin heavy chain, fast skeletal muscle-like | Keratin, type II cytoskeletal 5 |
| 3 | ACTC protein |  | TATA box binding protein associated factor (TAF) histone-like fold domain-containing protein | Keratin, type II cytoskeletal 5 | Phosphopyruvate hydratase | Collagen alpha-2(I) chain |
| 4 | Heat shock protein 70 |  | Elongation factor 1-alpha | 40S ribosomal protein S26 | Fast myotomal muscle tropomyosin (Tropomyosin alpha-1 chain) | Keratin, type II cytoskeletal 6A |
| 5 | Elongation factor 1-alpha |  | Myoglobin | Filamin-C | Fructose-bisphosphate aldolase | Glyceraldehyde-3-phosphate dehydrogenase |

#### Do all proteins have an equal chance of survival?

A central aim of this study was to investigate whether there is a bias towards or against the detection of certain food proteins. We sought to investigate if all proteins followed the same decay trend, ie. highly abundant in the fresh food, then reducing in abundance when cooked (and entrapped in foodcrust) and then further reducing in abundance when buried. To explore this, hierarchical cluster plots based on peptide abundance grouped by leading razor protein were created (Figure 4). For this, only proteins present in all three buried foodcrust replicates with >2 peptide matches were explored, as they were considered to consistently preserve at a quantity appreciable in general palaeoproteomic analysis. The cluster plots revealed that preservation varies by specific protein, and that not all proteins follow the same trend. One notable observation is an increased number of peptide matches to particular proteins in cooked samples when compared to the fresh ingredient, for example: Troponin T3, Fast Skeletal Type (TNNT3), myoglobin (MB) and globulin-family profile domain-containing protein (haemoglobin) in deer, and phosphopyruvate hydratase (enolase) and glyceraldehyde-3-phosphate dehydrogenase (GAPDH), and Fructose-bisphosphate aldolase in salmon (Figure 4). One explanation for this phenomenon could be the role played by heat in denaturing proteins, partly degrading them and opening them up so that enzymatic cleavage is more efficient. In the buried foodcrusts, while the overall number of proteins is lower than unburied samples, some proteins retained relatively high peptide abundance. In the deer samples these included globulin-family profile domain-containing protein (haemoglobin) and ACTC protein (actin) which were relatively abundant in all three buried foodcrust replicates, and in the salmon samples the comparatively abundant proteins included myosin heavy chain fast skeletal muscle-like, glyceraldehyde-3-phosphate dehydrogenase (GAPDH) and fast myotomal muscle tropomyosin. In contrast, while phosphopyruvate hydratase (enolase) is the most abundant protein in fresh and cooked deer replicates, it was found in relatively low abundance in buried foodcrust samples. This reveals that protein preservation is variable: certain proteins persist particularly well in buried foodcrust while others do not.

**Figure 4:**
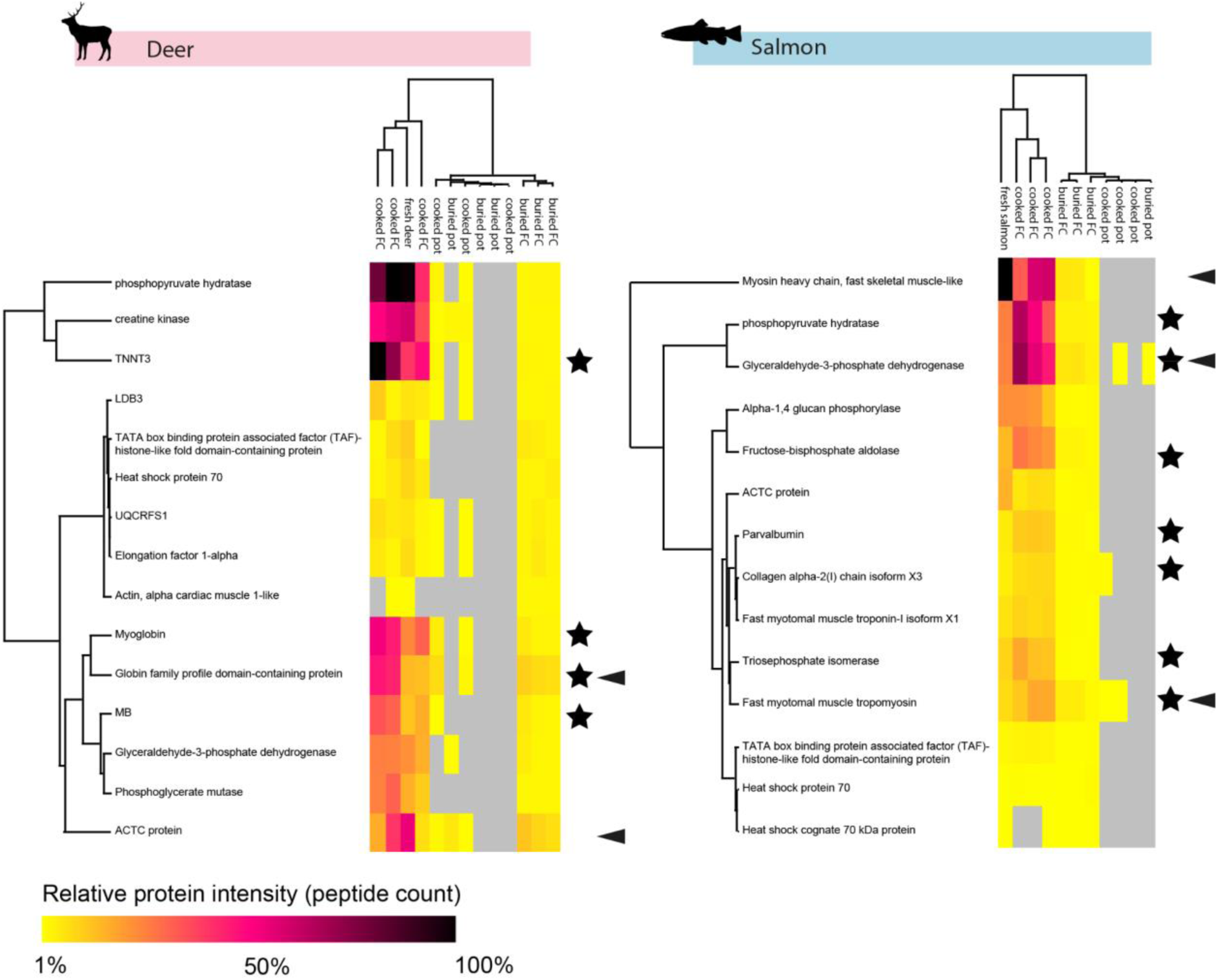
Heat map and dendrogram of proteins present in all buried deer (left) and salmon (right) foodcrust and ceramic replicates with peptide count >2. Euclidean clustering. Stars indicate proteins which are more abundant in cooked foodcrust samples than in the corresponding fresh ingredient sample. Arrows indicate proteins that are relatively abundant in buried foodcrusts compared to other proteins. Created in Perseus version 1.6.14.

#### Do buried foodcrust results reflect initial ingredient input?

The buried foodcrust samples were intended as analogues of archaeological foodcrusts, and thus to provide a baseline for the extractome that might be expected from archaeological foodcrusts. We investigated the extent to which buried cooked foodcrusts resemble the input protein composition. “Upset diagrams’’, an alternative to Venn diagrams [61], which were used to investigate the proteins shared by each sample type (Figure 3B). They revealed that for all ingredients, the fresh and cooked foodcrust samples contained the most common shared proteins of any set of samples, indicating that they are compositionally most similar. The largest overlapping group following this was fresh ingredients, cooked food crust and buried foodcrust in all cases, indicating that despite the reduction of proteins in buried samples, they still somewhat reflect the initial input composition.

To understand whether buried samples revealed the input ingredient taxonomy, Unipept Desktop was used to assign LCA for each peptide. Data were filtered following commonly used proteomic standards (greater than two PSMs to support a protein). In contrast to the buried ceramic samples which no harboured species-specific protein results, all buried deer and salmon foodcrust replicates produced sufficient proteomic evidence to identify the specific input ingredient to a species level in at least one replicate (supplementary text S3), while two of the chestnut replicates provided tissue specific evidence with some level of taxonomic specificity (Fagacea or *Quercus lobata*). Previously, a correlation between the presence of fish products and foodcrusts has been noted [24]. We also note that fish (and deer) proteins are more likely to be preserved in foodcrusts, but the presence of plants in foodcrust appears underrepresented in protein data. We particularly note that field cross-contamination from salmon and deer was more abundant in buried chestnut foodcrusts than were the input chestnut proteins (Table 2). Therefore proteomics may not be an appropriate single-method through which to address questions of plant processing in antiquity, at least by the methods adopted here. It is apparent that plants generate foodcrusts, but their molecular detection within foodcrusts remains challenging.

These results show that the input ingredient strongly influences the frequency of protein and peptide identifications in foodcrusts and ceramics. Chestnut proteins and peptides were identified less frequently than salmon or deer in buried foodcrusts, despite having higher protein and peptide abundance in fresh samples, and higher or similar protein and content to deer in cooked foodcrust (Figure 2). This leads us to believe that the comparably low preservation of chestnut in buried samples genuinely reflects their preservation potential relative to the other ingredients rather than other potential explanations such as chestnuts’ lower protein content or the fact that it underwent fewer cooking repeats, indicating an important bias in proteomic analysis of foodcrust residues. While in theory proteomics is capable of detecting proteins from any species represented in reference databases, in practice it appears that certain ingredients are more likely to be preserved or detected than others. Similarly, this has been observed in the analysis of ancient dental calculus, where a bias towards the detection of milk proteins over other dietary derived proteins has been reported [12]. This has obvious implications on the interpretation of archaeological results, for example, rendering plants less visible compared to other ingredients.

Furthermore, in this study, the lack of annotated proteins from some plant species has become starkly apparent. We note that a large number of the peptides identified in the chestnut samples matched to uncharacterised proteins, rendering their analysis difficult. Moreover, the absence of a *Castanea sativa* proteome in Uniprot at the time of analysis necessitated the use of *Castanea mollissima* in this investigation - which may impact identifications. This concurs with Hendy et al.’s previous comment on the dependence of shotgun proteomics on available databases, and its impact on plant identification [8]. Plants which have much larger proteomes are often absent, particularly for species that are not of current commercial relevance, such as heirloom cultivars. Database absence likely contributes to the lower detection rate of plants in archaeological samples or their detection at higher taxonomic specificity.

### Exploration of characteristics enabling protein survival in buried foodcrust samples

#### Why do particular proteins survive cooking and burial?

Having identified which proteins persist after cooking, foodcrust formation and burial, as well as the overall trend in the number of proteins preserved, we now explore whether these proteins harbour particular characteristics which may facilitate their survival. In this study it is apparent that degradation is not universal nor linear between different proteins [in contrast to 16]. As discussed above, certain proteins persist particularly well in buried samples while others do not (Figure 4), leading us to hypothesise that individual protein properties aid in their preservation. Previously, particular characteristics have been hypothesised to impact protein preservation in or on pottery and other mineral surfaces [14–16,52,62–64]. We wished to explore if there were particular characteristics on either a peptide or protein level that may be impacting the potential preservation of proteins in buried foodcrust samples.

A range of peptide and protein characteristics were investigated. These included concentration of different amino acids, peptide length, peptide hydropathicity, peptide isoelectric point, the sample’s lipid content, protein melting temperature, disorder prediction, amyloid propensity, protein secondary structure and relative solvent accessibility at a given peptide (Figure 5). The characteristics of bulk amino acid concentration, peptide length, peptide hydropathicity, and bulk deamidation were calculated manually, while various tools were used to calculate the other characteristics. These included: Protein thermal stability: DeepSTABp [65], Amyloid propensity: AMYPred-FRL[66], Disorder prediction: IUPred [67], Isoelectric point: IPC [68], and protein secondary structure and relative solvent accessibility: “Predict_Property”, a standalone, offline version of RaptorX web server [69,70](https://github.com/realbigws/Predict_Property). A script was written to extract the secondary structure, RSA, and disorder prediction for each peptide in the dataset (https://github.com/miranda-e/peptide_property_analyser). All characteristic results were then compiled and displayed using an R script (Figure 5). The full details of data analysis are present in supplementary text S4. While we initially aimed to explore the relationship between protein tertiary structure and preservation this proved challenging due to the paucity of tools for this analysis, and the poorly annotated or modelled nature of many target proteins. As an alternate way of investigating structural characteristics, we include secondary structure, disorder prediction and amyloid prediction. The characteristics of protein function and cell location were initially investigated through Gene Ontology terms, however these explorations were limited by relatively low levels of annotation in the target species’ databases (supplementary text S6) so this approach was not pursued. Factors other than cooking and burial such as the extraction protocol and data analysis parameters will have also impacted the composition of the extractome, however, all samples have experienced the same extraction and search protocol. Buried chestnut foodcrust data was removed from analysis due to very low peptide counts, and high levels of field cross-contamination, rendering any interpretation with statistical weight challenging.

**Figure 5:**
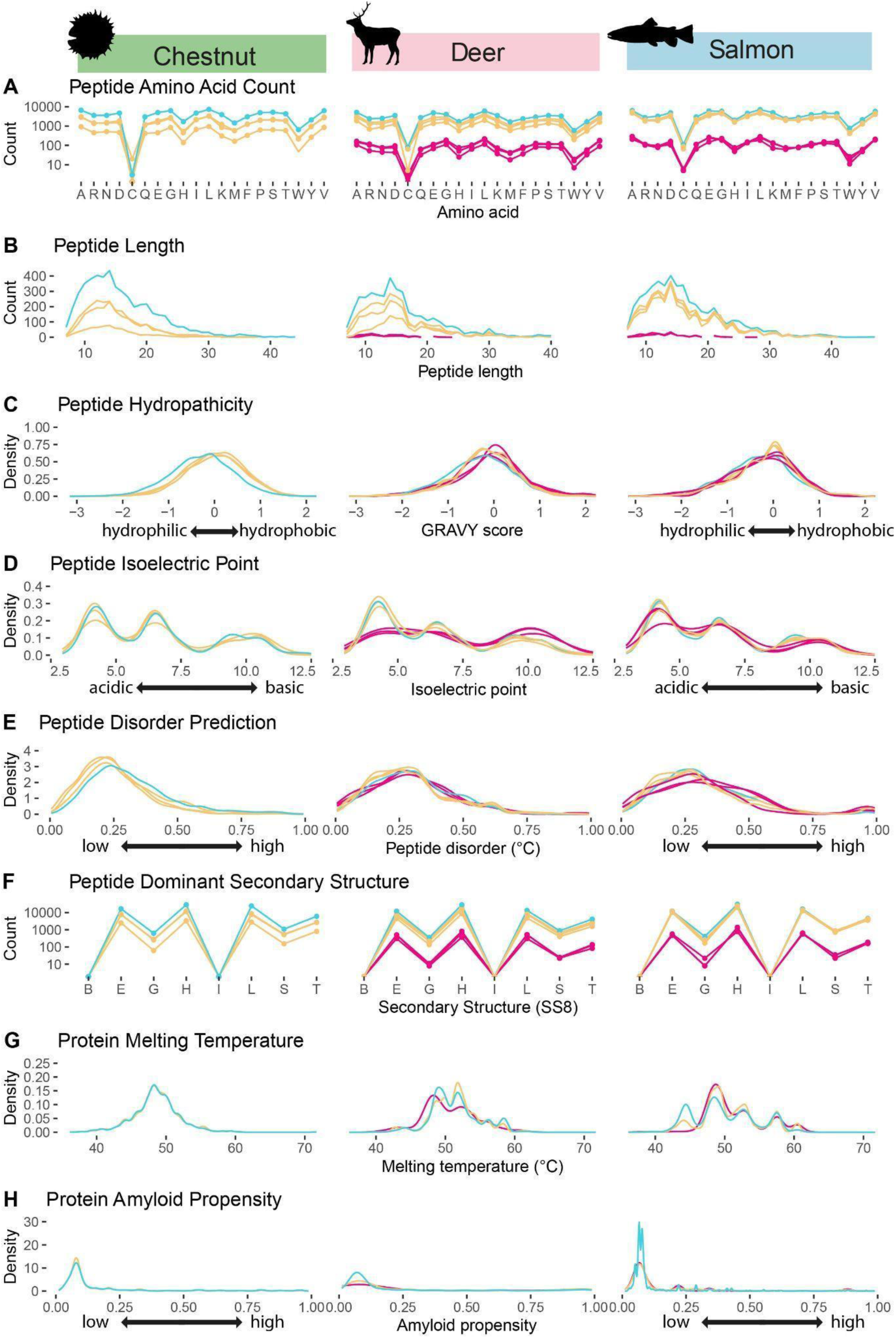
Protein and peptide characteristics for fresh, cooked foodcrust and buried foodcrust extractomes of chestnut, deer, and salmon, including: A: peptide amino acid count (points connected for visibility), B: Peptide Length, C: Peptide Hydropathicity, D: Peptide Isoelectric Point, E: Peptide Disorder Prediction, F: Peptide Dominant Secondary Structure (points connected for visibility), G: Protein Melting Temperature and H: Protein Amyloid Propensity. Buried chestnut replicates removed due to insufficient sample size and field contamination.

#### Proteins that survive in cooked and buried foodcrust do not have particular amino acid compositions or secondary structures

Previously, the impact of reactive amino acid content has been noted as a factor influencing protein survival [16], as has the impact of higher order structure and the location of a peptide within the structure, which may protect or expose particular peptides [16,63]. Amino acid sequence also has been reported as a driving factor in protein abundance by determining conformational stability and reducing synthesis cost [71]. Bulk amino acid count (Figure 5a) and peptide secondary structure (Figure 5f) demonstrated no global change between their fresh, cooked and buried state for any ingredient, indicating that they likely did not play a substantial global role in peptide preservation. Secondary structure is innately linked to a protein’s function and stability, with certain structures being more stable. Secondary structure was collected using the following categories: G: 310 helix, H: alpha-helix, I: pi-helix, E: beta-strand, B:beta-bridge, T: beta-turn, S: high curvature loop, and L for irregular (Figure 5f). Secondary structures have been reported to be distributed across all proteins in the following ratio; alpha-helix, beta-strand, irregular, beta-turn, high curvature loop,310 helix, beta-bridge, pi-helix = 34:21:20:11:9:4:1:0 [72]. Similar ratios inline with the background distribution were observed in all samples, with only minor variation between input ingredients - meaning that secondary structure does not seem to be a factor in determining which peptides we detect in the extractome. Moreover, as secondary structure and bulk amino acid distribution were no different in cooked or buried samples for any ingredient, it appears that these characteristics do not impact protein preservation.

#### More hydrophobic peptides are slightly more likely to survive cooking and burial

Previously, the potential role of protein hydropathicity in protein preservation has been hypothesised, whereby hydrophilic proteins leach from ceramics during washing and/or burial [15,16,52]. In this study we investigated whether peptide hydropathicity (solubility) correlated with peptide abundance in fresh, cooked or buried samples, with the hypothesis that less water soluble (hydrophobic) peptides might preferentially survive in buried samples. Grand Average of Hydropathy (GRAVY) score, a standard measure of protein polarity, was calculated on a peptide level for fresh, cooked food crust and buried foodcrust samples (Figure 5c). For both salmon and deer, GRAVY score increased slightly in cooked and buried foodcrusts compared to fresh samples, indicating that peptides were generally slightly more hydrophobic in cooked and buried samples than in fresh samples. This result supports previous hypotheses that water leaching over time may reduce soluble protein content in pottery [16,52], leaving slightly more hydrophobic proteins to be detected in higher abundance in buried samples, although we note that a slight increase in hydrophobic peptides was also observed after cooking, indicating that hydrophilic peptides may also be less likely to be entrapped in foodcrust.

#### Cooking may liberate peptides located deep within the protein’s 3D structure

The potential role of higher order structure and the location of a peptide within the structure, which may protect or expose particular peptides has been noted [17,72]. Relative solvent accessibility (RSA) is a measure of the exposure of an amino acid within its tertiary structure, and therefore how accessible that residue is to solvents (ie. amino acids located deeper within the 3D structure are less accessible to solvents). The results revealed that cooked foodcrusts had a higher proportion of peptides with deep RSA (ie. peptides with amino acids located deep within the tertiary protein structure), than did fresh ingredients, which is most marked in chestnut. One explanation for this is that as tertiary protein structures unfold during denaturation during cooking, peptides which are located deep within the protein structure become more accessible to extraction than they are in uncooked ingredients. This means that in uncooked archaeological samples we are probably less likely to see deep peptides than in their cooked counterparts. In buried foodcrust samples the proportion of “deep” RSA peptides continued to decrease in deer samples but increased in salmon samples, providing inconsistent results.

#### Protein thermal stability may impact peptide survival

Thermal stability is the ability of proteins to resist changes in structure caused by heating. We investigated this characteristic with the hypothesis that thermostable peptides would persist through the cooking, entrapment, and burial process. Melting temperature (Tm) is often used as a measure of protein thermal stability. Proteins surviving in buried salmon foodcrust are slightly more thermostable than the fresh ingredient and unburied foodcrust, with fewer peptides from proteins of low thermal stability (Tm < 45°C) surviving in buried foodcrust replicates (Figure 5g). This demonstrates that thermally stable peptides were more likely to survive in buried salmon than proteins with lower thermal stability. Conversely in buried deer foodcrust, peptides from less thermally stable proteins survived better than peptides with higher thermal stability, further demonstrating the varied behaviour of different ingredients.

#### Certain properties may aid in the preservation of particular ingredients or proteins

Some characteristics demonstrated changes after cooking and burial only for particular ingredients. Amyloid propensity and disorder prediction showed changes primarily for salmon, but not in deer samples. Previously Collins et al. [64] speculated that entropic effects would promote survival of flexible molecules that could adapt and bind to the mineral surface. Demarchi et al. provided evidence that mineral binding of a small flexible acid rich region was responsible for the persistence of a peptide form of a c-type lectin of African ostrich eggshell into deep time [73]. Most recently, Scott [74] proposed that the robust nature of amyloid fibrils and other factors contributing to protein aggregation may explain the presence of particular proteins and peptides in the archaeological record, noting that dietary proteins persisting in ancient dental calculus are often amyloidogenic.The analysis of intrinsically disordered proteins revealed that in the case of salmon, the proportion of peptides with high disorder prediction slightly increased following cooking and remained high during burial meaning that more flexible peptides become relatively more representative than inflexible ones (Figure 5e). Deer samples did not display this trend. Similarly, the analysis of amyloid propensity revealed that peptides which could readily stack were more likely to be detected in buried salmon samples than fresh or cooked (Figure 5h), potentially indicating that this characteristic plays a role in the preservation of some salmon proteins. This trend was also not observed in deer samples. In the case of deer, the density of peptides with low isoelectric points (i.e. acidic, water soluble peptides) decreased in the buried samples compared to the fresh ingredient and unburied foodcrust, while the density of peptides with high pIs (basic peptides) increased (Figure 5d), demonstrating that acidic peptides survived poorly in buried deer foodcrust, a change not observed in salmon or chestnut. This indicates that certain characteristics may aid in the preservation of particular ingredients or proteins.

While the buried chestnut samples were not included in the broader characterisation analysis due to small sample size, we note that the only two chestnut-specific proteins to survive in buried foodcrust samples (Cupin type-1 domain-containing protein and Chitin-binding type-1 domain-containing protein) are both allergenic. Allergenic proteins are often characterised by their stability, either in terms of their tertiary structure (such as the beta-barrel observed in Cupin-type proteins), or resistance to heat or digestive degradation [75], and have previously been observed to preferentially preserve in ancient dental calculus [76].

The potential role of lipids in creating water-impermeable barriers which may shelter proteins in pottery from forces of degradation has previously been noted [17], although lipids may complicate protein extraction [15]. A vast body of work has explored the impact of organic content on protein and nitrogen preservation in sediments [77–82]. Previously, some of the buried foodcrust replicates were analysed by flame ionisation detector [47], facilitating an opportunity to examine any correlations between protein and lipid content in the same vessel.The total peptide and protein count was compared to these lipid quantities generated by Bondetti et al. [47] for each sample. We note that there was a surprising level of variation in lipid concentration between replicates of identical input ingredients and weight. This revealed that in addition to the impact of cooking practice and frequency of use [50], even identically processed foodcrusts are not homogenous. However due to the small sample size it was not possible to reveal the impact of lipid content on protein detection (supplementary text S5). Future controlled dosing studies would further address the impact of lipid content on protein preservations in ceramics and their residues.

#### Markers of diagenesis

Peptide length was investigated as a potential indicator of diagenesis from the cooking and entrapment process and/or from burial. Differences in peptide length were observed between fresh ingredients, cooked foodcrust and buried foodcrust. For all ingredients, the distribution indicated that shorter peptides were more likely to be detected in all samples (Figure 5b). Peptide length decreased upon cooking, with chestnut providing the most marked reduction, and salmon the least. It is notable that long peptides (above ∼ 20 amino acids) were very rarely detected in buried samples of any ingredient. This may demonstrate that long peptides are rarely preserved in buried samples, or alternatively, as long peptides are always less abundant, that the smaller sample size of buried peptides reduces the probability of their detection. As peptide length decreases, it seems likely that the probability of identifying a tissue and taxonomically-specific sequence would also decrease. This analysis further revealed differences in the behaviour of the different ingredients.

Deamidation of asparagine and glutamine was explored as a marker of preservational quality [83]. In certain proteins deamidation has been reported to change protein tertiary structure [84]. In the case of deer and salmon, the highest proportion of peptides bearing deamidation were the unburied foodcrust samples, demonstrating that heating plays a role in deamidation. In the case of salmon and deer, the proportion of deamidated peptides fell after burial (Figure 6). Notably, the proportion of deamidated peptides does not continue to increase in the buried state considering that deamidation has been used as an indicator of protein preservation quality, perhaps because the majority of possible deamidations have already occurred during heating, and timescales of chemically mediated deamidation are slow at burial temperatures [85]. This demonstrates that deamidation may not be a good indicator of peptide authenticity where modern contaminants may have undergone heating. Unfortunately, due to the necessary peptide filtration steps discussed below, it was not possible to apply deamiDATE here to assess site-specific deamidation. Further investigation of site-specific PTMs may reveal evidence for aspects of food preparation and taphonomy.

**Figure 6:**
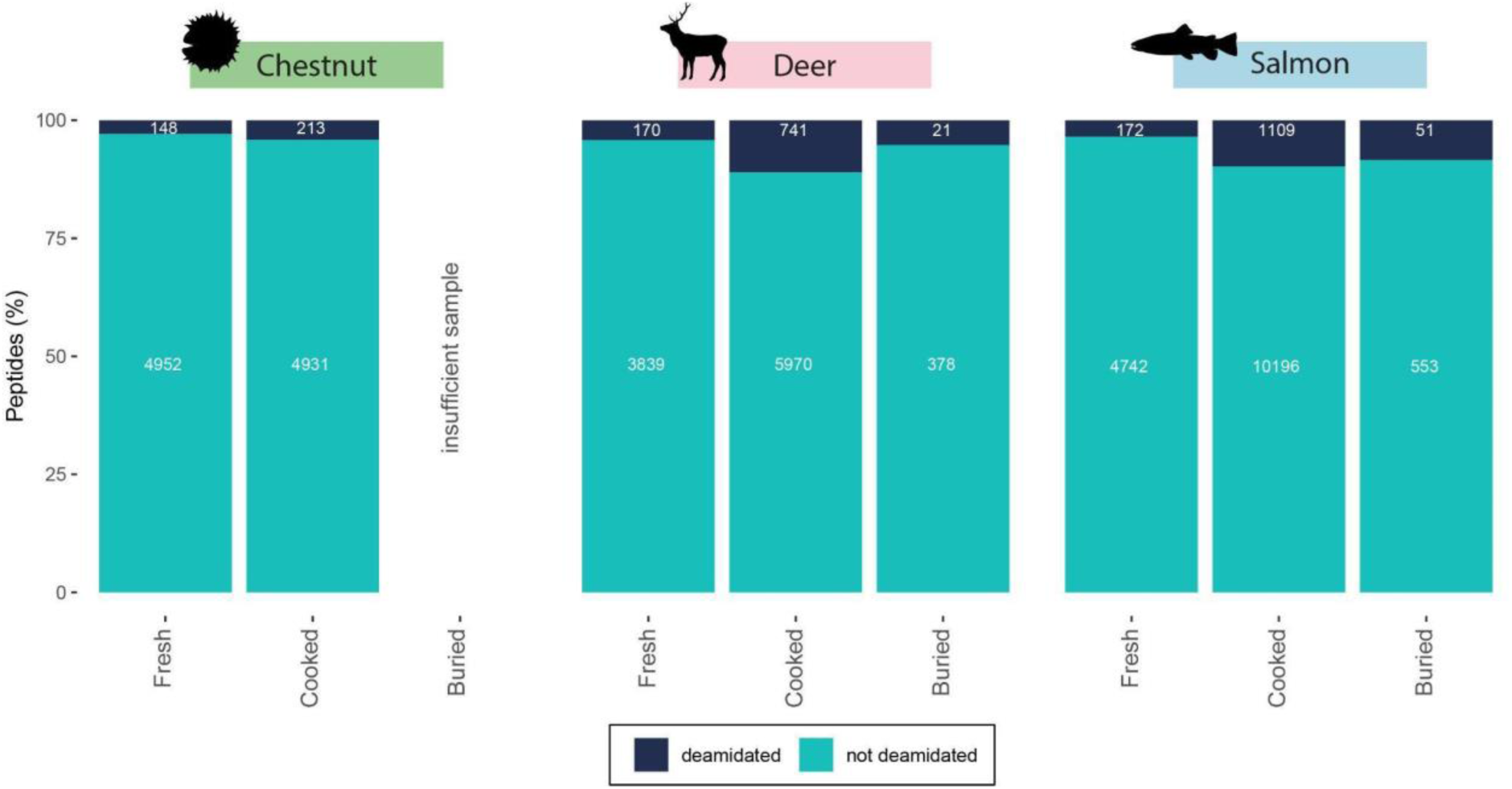
Proportion of deamidated peptides. Note: cooked and buried foodcrust replicates =3, Fresh ingredient replicates =1.

#### Case study: haemoglobin

While no single physiochemical characteristic investigated appears to explain why some proteins survive better than others in buried foodcrusts, it is apparent that these properties may play a role for individual ingredients and proteins. One protein that preserves relatively well in the foodcrust samples is globin-domain containing protein (haemoglobin), so we investigated this protein in greater detail as a case study to understand, on an individual protein level, if particular characteristics may be aiding in its survival. These are likely too complex and varied to be detected broadly, across the whole sample set.

GRAVY score, amyloid propensity, secondary structure and RSA were overlaid on top of a peptide abundance heatmap (Figure 7), illustrating that in the case of the globin family protein, the peptides which preserve most commonly in the buried sample are located on the most hydrophobic parts of the protein, and often in regions with high amyloid propensity (Figure 7). This demonstrates in the case of globin that hydrophobic (insoluble) peptides appear to be more likely to persist in our buried samples than hydrophilic peptides. However this means that a complex array of mechanisms, interactions and protein compositions must be impacting the survival of proteins and peptides in buried foodcrusts.

**Figure 7:**
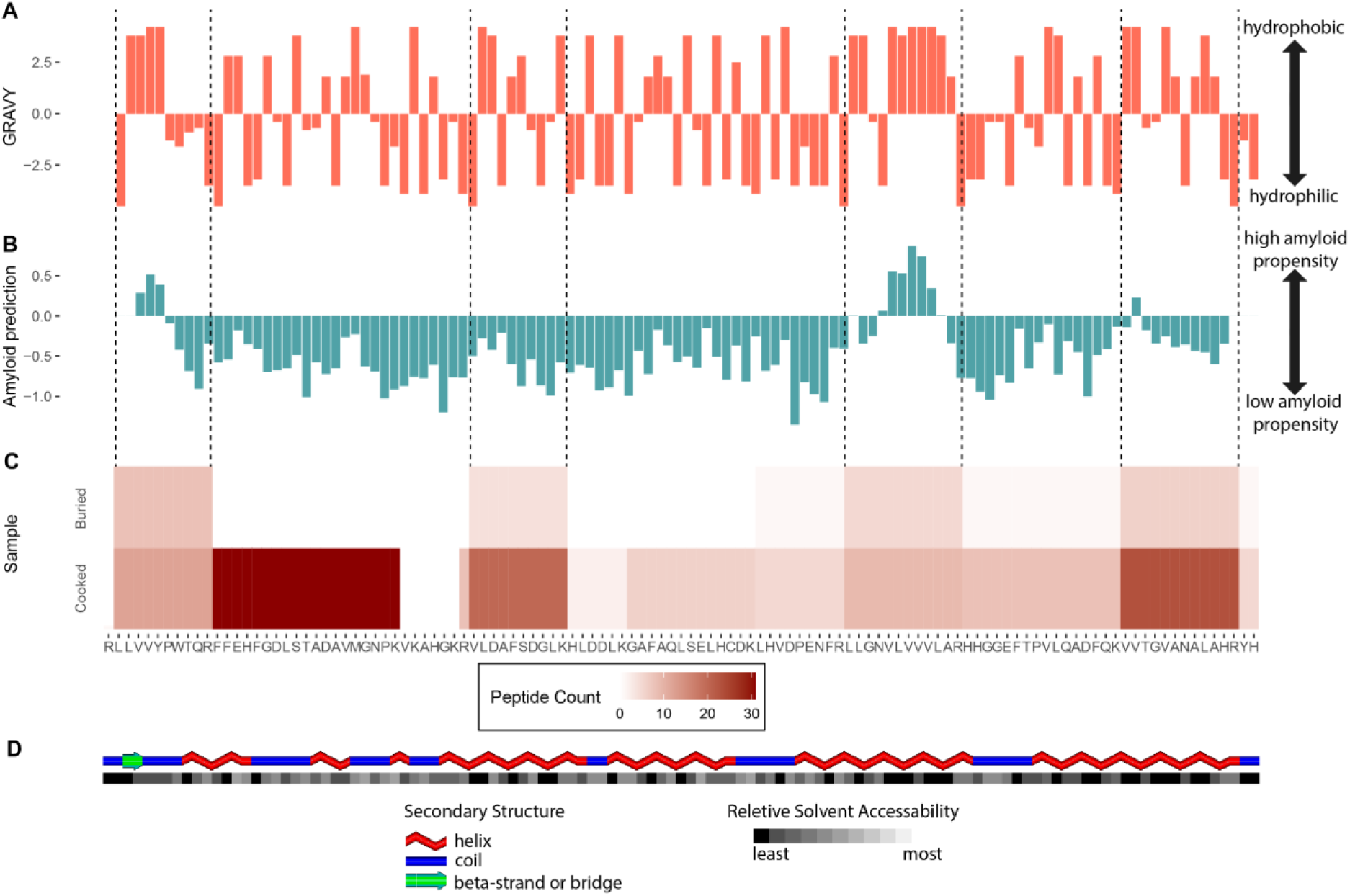
Globin characteristic: hydropathicity (A), amyloid prediction (B) with heatmaps of peptide counts for each region (C) of buried (top) and cooked (bottom) foodcrust deer samples, (D) secondary structure of sequence: blue= coil, red= helix and relative solvent accessibility (RSA) Black=completely buried, White=fully exposed (created in POLYVIEW-2D [86]).

#### No property single-handedly explains why particular proteins survive cooking and burial

In this study, many protein characteristics were investigated to understand potential explanations for why certain proteins were detected in buried foodcrusts. We observe that the input ingredient was the most influential factor in protein survival, and individual proteins follow different preservation trends. For example, in the foodcrust created by cooking deer meat, basic peptides seem to preferentially survive the cooking and burial environment. But this trend was not observed for proteins detected from salmon foodcrust. Similarly, proteins with higher thermal stability, disorder prediction, and amyloid propensity were preferentially preserved in buried salmon foodcrusts, yet not in deer. Some trends that were observed across all food types including that slightly hydrophobic peptides are more likely to preserve after cooking and burial, that the recovery of longer peptides tended to decrease after cooking and burial, and that deamidation increased following cooking, but not necessarily burial. However no trend could single handedly explain why particular peptides and proteins survive cooking and burial. While few trends in protein characteristic were observed across all ingredients, when viewed on an individual protein level, haemoglobin peptides that were slightly hydrophobic and had high amyloid propensity were more likely to preserve in buried foodcrust samples, even though they were not particularly abundant in the unburied sample. This indicates that particular protein or peptide characteristics are indeed involved in determining which proteins/peptides survive in buried samples, but that these complex interactions are likely to vary based on a myriad of factors, and may be obscured when the data is viewed more broadly.

### Future directions

A key goal of this study is to consider expectations for the survival of proteins in foodcrusts and ceramics in archaeological contexts. While the buried foodcrusts in this investigation revealed a number of protein identifications with an informative level of protein and taxa-specificity, it is important to note that the samples were buried for only 6 months and in a temperate climate, and therefore may not be comparable to samples of much older temporal and thermal age. While Barker et al. [16] reported that protein content dropped rapidly upon burial and at a slower rate thereafter, the exact rate of dietary protein decay over longer periods of time and in other climatic conditions is poorly understood. The results of this study should therefore be seen as an upper limit to protein preservation, as it may well be that samples of considerable age or from hot climates may not yield confidently preserved proteins, necessitating a cautious approach to the destruction of valuable samples.

In this experiment the input ingredient itself appeared to play a strong role in the difference in protein preservation/detection between samples. This has implications on ingredient visibility in archaeological interpretations on foodcrust results, particularly for plants. Future investigation of the impact of ingredient mixing on protein preservation will be necessary to understand the full implications of this. Furthermore, further investigation into the impact of protein interactions with other macronutrients such as lipids and carbohydrates on protein preservation is required to fully understand this issue.

We are aware that extraction and LC-MS/MS protocols will have impacted the peptides which are detected. As a result, we are viewing the detected proteins and peptides through a lens of the analytical processes they have been through. Some of these may be taphonomic or cultural such as cooking and burial, but others are inflicted by post depositional processes, storage, handling and extraction. Untangling the role of each will be imperative in understanding the impact of any one variable. The extraction of proteins from ceramics continues to be a challenge, with ongoing optimisation [14–16,21,51,63,87,88], which will undoubtedly impact protein extraction.

## Conclusion

In this investigation we sought to explore the utility of proteomic analysis of foodcrusts and ceramics in understanding ancient food preparation, by examining the extent to which the buried foodcrust extractome reflects input ingredients. The results revealed that foodcrusts harboured more preserved proteins than ceramics, and we note, in line with previous studies, that proteins are not easily extracted from ceramic matrices [14,15]. Sufficient taxonomic and tissue-specific identifications were made to detect the relevant ingredient in buried foodcrust created by cooking deer and salmon meat, and sometimes chestnut flour, demonstrating that ancient foodcrusts may be a viable matrix from which to extract dietary proteins, but that not all ingredients were equal in protein retrieval revealing that input ingredient biases protein recovery. Preservation was not universal between proteins and peptides; those which were most abundant in the fresh meat and flour were not necessarily the most frequently identified in buried samples. These biases have implications on archaeological interpretations of ancient foodcrusts and ceramics, namely that certain ingredients and proteins will be unlikely to be detected in ancient samples. It is clear further work is necessary to understand the biases of input ingredients, particularly when they are mixed, and over longer periods of vessel use and burial. Secondly we attempted to characterise the proteins and peptides retrieved from buried foodcrust samples, to understand physicochemical factors influencing their preservation. While we observed that more hydrophobic peptides were slightly more likely to survive cooking and burial, no property was seen to single-handedly explain why particular proteins/peptides survive in buried foodcrusts, with results indicating that certain properties act on protein preservation in complex ways requiring further investigation, or that characteristics not investigated here may play a role. This study demonstrates the value of experimental analyses to anticipate a maximum baseline of protein results from archaeological samples.

## Data Availability

The mass spectrometry proteomics data have been deposited to the ProteomeXchange Consortium via the PRIDE [89] partner repository with the dataset identifier PXD050001 and 10.6019/PXD050001.

## Acknowledgements

We gratefully acknowledge the use of the Thermo Scientific Orbitrap Fusion™ Tribrid™ in the York Centre of Excellence in Mass Spectrometry. The centre was created thanks to a major capital investment through Science City York, supported by Yorkshire Forward with funds from the Northern Way Initiative, and subsequent support from EPSRC (EP/K039660/1; EP/M028127/1). We thank Niklas Hausmann for assistance with R, and Alexandre Lucquin for assistance locating lipid data files.

## AI-assisted technology declaration

AI-assisted technology was used to support code writing and debugging of the "peptide_property_analyser.py" python script created for this manuscript. The script was then checked by a software developer and tested extensively with test datasets of known output to validate it. AI-assisted technologies were not used in any other capacity in this manuscript.

## Works Cited

1. Runge AKW et al. 2021 Palaeoproteomic analyses of dog palaeofaeces reveal a preserved dietary and host digestive proteome. Proc. Biol. Sci. 288, 20210020. (doi:10.1098/rspb.2021.0020)

2. Colonese AC et al. 2017 New criteria for the molecular identification of cereal grains associated with archaeological artefacts. Sci. Rep. 7, 6633. (doi:10.1038/s41598-017-06390-x)

3. Yang Y, Shevchenko A, Knaust A, Abuduresule I, Li W, Hu X, Wang C, Shevchenko A. 2014 Proteomics evidence for kefir dairy in Early Bronze Age China. J. Archaeol. Sci. 45, 178–186. (doi:10.1016/j.jas.2014.02.005)

4. Hong C, Jiang H, Lü E, Wu Y, Guo L, Xie Y, Wang C, Yang Y. 2012 Identification of milk component in ancient food residue by proteomics. PLoS One 7, e37053. (doi:10.1371/journal.pone.0037053)

5. Xie M, Shevchenko A, Wang B, Shevchenko A, Wang C, Yang Y. 2016 Identification of a dairy product in the grass woven basket from Gumugou Cemetery (3800 BP, northwestern China). Quat. Int. 426, 158–165. (doi:10.1016/j.quaint.2016.04.015)

6. Buckley M, Melton ND, Montgomery J. 2013 Proteomics analysis of ancient food vessel stitching reveals >4000-year-old milk protein. Rapid Commun. Mass Spectrom. 27, 531–538. (doi:10.1002/rcm.6481)

7. Cappellini E et al. 2010 A multidisciplinary study of archaeological grape seeds. Naturwissenschaften 97, 205–217. (doi:10.1007/s00114-009-0629-3)

8. Hendy J et al. 2018 Ancient proteins from ceramic vessels at Çatalhöyük West reveal the hidden cuisine of early farmers. Nat. Commun. 9, 1–10. (doi:10.1038/s41467-018-06335-6)

9. Evans M et al. 2023 Detection of dairy products from multiple taxa in Late Neolithic pottery from Poland: an integrated biomolecular approach. Royal Society Open Science 10, 230124. (doi:10.1098/rsos.230124)

10. Jeong C et al. 2018 Bronze Age population dynamics and the rise of dairy pastoralism on the eastern Eurasian steppe. Proceedings of the National Academy of Sciences 115, E11248–E11255. (doi:10.1073/pnas.1813608115)

11. Wilkin S et al. 2021 Dairying enabled Early Bronze Age Yamnaya steppe expansions. Nature 598, 629–633. (doi:10.1038/s41586-021-03798-4)

12. Hendy J et al. 2018 Proteomic evidence of dietary sources in ancient dental calculus. Proc. Biol. Sci. 285, 20180977. (doi:10.1098/rspb.2018.0977)

13. Geber J, Tromp M, Scott A, Bouwman A, Nanni P, Grossmann J, Hendy J, Warinner C. 2019 Relief food subsistence revealed by microparticle and proteomic analyses of dental calculus from victims of the Great Irish Famine. Proc. Natl. Acad. Sci. U. S. A. 116, 19380–19385. (doi:10.1073/pnas.1908839116)

14. Craig OE, Collins MJ. 2002 The Removal of Protein from Mineral Surfaces: Implications for Residue Analysis of Archaeological Materials. J. Archaeol. Sci. 29, 1077–1082. (doi:10.1006/jasc.2001.0757)

15. Craig OE, Collins MJ. 2000 An improved method for the immunological detection of mineral bound protein using hydrofluoric acid and direct capture. J. Immunol. Methods 236, 89–97. (doi:10.1016/s0022-1759(99)00242-2)

16. Barker A, Dombrosky J, Venables B, Wolverton S. 2018 Taphonomy and negative results: An integrated approach to ceramic-bound protein residue analysis. J. Archaeol. Sci. 94, 32–43. (doi:10.1016/j.jas.2018.03.004)

17. Tanasi D, Cucina A, Cunsolo V, Saletti R, Di Francesco A, Greco E, Foti S. 2021 Paleoproteomic profiling of organic residues on prehistoric pottery from Malta. Amino Acids 53, 295–312. (doi:10.1007/s00726-021-02946-4)

18. Solazzo C, Fitzhugh WW, Rolando C, Tokarski C. 2008 Identification of Protein Remains in Archaeological Potsherds by Proteomics. Anal. Chem. 80, 4590– 4597. (doi:10.1021/ac800515v)

19. Chowdhury MP, Campbell S, Buckley M. 2021 Proteomic analysis of archaeological ceramics from Tell Khaiber, southern Iraq. J. Archaeol. Sci. 132, 105414. (doi:10.1016/j.jas.2021.105414)

20. Dallongeville S, Garnier N, Casasola DB, Bonifay M, Rolando C, Tokarski C. 2011 Dealing with the identification of protein species in ancient amphorae. Anal. Bioanal. Chem. 399, 3053–3063. (doi:10.1007/s00216-010-4218-2)

21. Pal Chowdhury M, Makarewicz C, Piezonka H, Buckley M. 2022 Novel Deep Eutectic Solvent-Based Protein Extraction Method for Pottery Residues and Archeological Implications. J. Proteome Res. 21, 2619–2634. (doi:10.1021/acs.jproteome.2c00340)

22. Admiraal M et al. 2023 The role of salmon fishing in the adoption of pottery technology in subarctic Alaska. J. Archaeol. Sci. 157, 105824. (doi:10.1016/j.jas.2023.105824)

23. Heron C, Craig OE. 2015 Aquatic Resources in Foodcrusts: Identification and Implication. Radiocarbon 57, 707–719. (doi:10.2458/azu_rc.57.18454)

24. Courel B et al. 2021 The use of early pottery by hunter-gatherers of the Eastern European forest-steppe. Quat. Sci. Rev. 269, 107143. (doi:10.1016/j.quascirev.2021.107143)

25. Lucquin A et al. 2016 Ancient lipids document continuity in the use of early hunter–gatherer pottery through 9,000 years of Japanese prehistory. Proceedings of the National Academy of Sciences 113, 3991–3996. (doi:10.1073/pnas.1522908113)

26. Craig OE. 2004 Organic analysis of «food crusts» from sites in the Schelde valley, Belgium: a preliminary evaluation. Notae Praehistoricae 24, 209–217.

27. Heron C, Craig OE, Luquin A, Steele VJ, Thompson A, Piličiauskas G. 2015 Cooking fish and drinking milk? Patterns in pottery use in the southeastern Baltic, 3300–2400 cal BC. J. Archaeol. Sci. 63, 33–43. (doi:10.1016/j.jas.2015.08.002)

28. Gunnarssone A, Oras E, Talbot HM, Ilves K, Legzdiņa D. 2020 Cooking for the living and the dead: Lipid analyses of rauši settlement and cemetery pottery from the 11th–13th century. Estonian J. Archaeol. 24, 45–69. (doi:10.3176/arch.2020.1.02)

29. Bondetti M et al. 2021 Neolithic farmers or Neolithic foragers? Organic residue analysis of early pottery from Rakushechny Yar on the Lower Don (Russia). Archaeol. Anthropol. Sci. 13, 141. (doi:10.1007/s12520-021-01412-2)

30. Gibbs K et al. 2017 Exploring the emergence of an ‘Aquatic’ Neolithic in the Russian Far East: organic residue analysis of early hunter-gatherer pottery from Sakhalin Island. Antiquity 91, 1484–1500. (doi:10.15184/aqy.2017.183)

31. Bondetti M, Scott S, Lucquin A, Meadows J, Lozovskaya O, Dolbunova E, Jordan P, Craig OE. 2020 Fruits, fish and the introduction of pottery in the Eastern European plain: Lipid residue analysis of ceramic vessels from Zamostje 2. Quat. Int. 541, 104–114. (doi:10.1016/j.quaint.2019.05.008)

32. Lucquin A et al. 2023 The impact of farming on prehistoric culinary practices throughout Northern Europe. Proc. Natl. Acad. Sci. U. S. A. 120, e2310138120. (doi:10.1073/pnas.2310138120)

33. Taché K, Jaffe Y, Craig OE, Lucquin A, Zhou J, Wang H, Jiang S, Standall E, Flad RK. 2021 What do ‘barbarians’ eat? Integrating ceramic use-wear and residue analysis in the study of food and society at the margins of Bronze Age China. PLoS One 16, e0250819. (doi:10.1371/journal.pone.0250819)

34. Shoda S, Lucquin A, Sou CI, Nishida Y, Sun G, Kitano H, Son J-H, Nakamura S, Craig OE. 2018 Molecular and isotopic evidence for the processing of starchy plants in Early Neolithic pottery from China. Sci. Rep. 8, 17044. (doi:10.1038/s41598-018-35227-4)

35. Yoneda M, Kisida K, Gakuhari T, Omori T, Abe Y. 2019 Interpretation of bulk nitrogen and carbon isotopes in archaeological foodcrusts on potsherds. Rapid Commun. Mass Spectrom. 33, 1097–1106. (doi:10.1002/rcm.8446)

36. Craig OE et al. 2013 Earliest evidence for the use of pottery. Nature 496, 351–354. (doi:10.1038/nature12109)

37. Shoda S, Lucquin A, Ahn J-H, Hwang C-J, Craig OE. 2017 Pottery use by early Holocene hunter-gatherers of the Korean peninsula closely linked with the exploitation of marine resources. Quat. Sci. Rev. 170, 164–173. (doi:10.1016/j.quascirev.2017.06.032)

38. Admiraal M et al. 2023 Chemical analysis of pottery reveals the transition from a maritime to a plant-based economy in pre-colonial coastal Brazil. Sci. Rep. 13, 16771. (doi:10.1038/s41598-023-42662-5)

39. Robson HK et al. 2022 Light Production by Ceramic Using Hunter-Gatherer-Fishers of the Circum-Baltic. Proceedings of the Prehistoric Society 88, 25–52. (doi:10.1017/ppr.2022.12)

40. Oras E, Lucquin A, Lõugas L, Tõrv M, Kriiska A, Craig OE. 2017 The adoption of pottery by north-east European hunter-gatherers: Evidence from lipid residue analysis. J. Archaeol. Sci. 78, 112–119. (doi:10.1016/j.jas.2016.11.010)

41. Regert M, Vacher S, Moulherat C, Decavallas O. 2003 Adhesive production and pottery function during the iron age at the site of grand aunay (sarthe, France). Archaeometry 45, 101–120. (doi:10.1111/1475-4754.00098)

42. Lyu N, Yan L, Wang T, Lin L, Rao H, Yang Y. 2024 Characterization of Pottery Foodcrusts Through Lipid and Proteomic Analyses: A Case Study from the Xiawan Site in Yixing City, East China. J. Archaeol. Sci., 161, 105902. (doi:10.1016/j.jas.2023.105902)

43. Shevchenko A, Schuhmann A, Thomas H, Wetzel G. 2018 Fine Endmesolithic fish caviar meal discovered by proteomics in foodcrusts from archaeological site Friesack 4 (Brandenburg, Germany). PLoS One 13, e0206483. (doi:10.1371/journal.pone.0206483)

44. Miller MJ et al. 2020 Interpreting ancient food practices: stable isotope and molecular analyses of visible and absorbed residues from a year-long cooking experiment. Sci. Rep. 10, 1–16. (doi:10.1038/s41598-020-70109-8)

45. Evershed RP, Copley MS, Dickson L, Hansel FA. 2008 Experimental evidence for the processing of marine animal products and other commodities containing polyunsaturated fatty acids in pottery vessels. Archaeometry 50, 101–113. (doi:10.1111/j.1475-4754.2007.00368.x)

46. Charters S, Evershed RP, Quye A, Blinkhorn PW, Reeves V. 1997 Simulation Experiments for Determining the Use of Ancient Pottery Vessels: the Behaviour of Epicuticular Leaf Wax During Boiling of a Leafy Vegetable. J. Archaeol. Sci. 24, 1–7. (doi:10.1006/jasc.1995.0091)

47. Bondetti M, Scott E, Courel B, Lucquin A, Shoda S, Lundy J, Labra-Odde C, Drieu L, Craig OE. 2021 Investigating the formation and diagnostic value of ω -( o - alkylphenyl)alkanoic acids in ancient pottery. Archaeometry 63, 594–608. (doi:10.1111/arcm.12631)

48. Liu X, Ren M, Fu Y, Hu Y, Wang S, Yang Y. 2023 New insights into the use of Neolithic pottery in Guangxi of South China: organic residue analysis of experimental and archaeological pottery. Heritage Science 11, 1–15. (doi:10.1186/s40494-023-01045-9)

49. Hammann S, Cramp LJE. 2018 Towards the detection of dietary cereal processing through absorbed lipid biomarkers in archaeological pottery. J. Archaeol. Sci. 93, 74–81. (doi:10.1016/j.jas.2018.02.017)

50. Drieu L, Regert M, Mazuy A, Vieugué J, Bocoum H, Mayor A. 2022 Relationships between lipid profiles and use of ethnographic pottery: An exploratory study. J. Archaeol. Method Theory 29, 1294–1322. (doi:10.1007/s10816-021-09547-1)

51. Barker A, Venables B, Stevens SM, Seeley KW, Wang P, Wolverton S. 2012 An Optimized Approach for Protein Residue Extraction and Identification from Ceramics After Cooking. Journal of Archaeological Method and Theory 19, 407–439.

52. Evershed RP, Tuross N. 1996 Proteinaceous Material from Potsherds and Associated Soils. J. Archaeol. Sci. 23, 429–436. (doi:10.1006/jasc.1996.0038)

53. Hughes CS, Foehr S, Garfield DA, Furlong EE, Steinmetz LM, Krijgsveld J. 2014 Ultrasensitive proteome analysis using paramagnetic bead technology. Mol. Syst. Biol. 10, 757. (doi:10.15252/msb.20145625)

54. Hughes CS, Moggridge S, Müller T, Sorensen PH, Morin GB, Krijgsveld J. 2018 Single-pot, solid-phase-enhanced sample preparation for proteomics experiments. Nat. Protoc. 14, 68–85. (doi:10.1038/s41596-018-0082-x)

55. Cleland TP. 2018 Human Bone Paleoproteomics Utilizing the Single-Pot, Solid-Phase-Enhanced Sample Preparation Method to Maximize Detected Proteins and Reduce Humics. J. Proteome Res. 17, 3976–3983. (doi:10.1021/acs.jproteome.8b00637)

56. Bleasdale M et al. 2021 Ancient proteins provide evidence of dairy consumption in eastern Africa. Nat. Commun. 12, 1–11. (doi:10.1038/s41467-020-20682-3)

57. Ventresca Miller AR et al. 2022 The spread of herds and horses into the Altai: How livestock and dairying drove social complexity in Mongolia. PLoS One 17, e0265775. (doi:10.1371/journal.pone.0265775)

58. Sakalauskaite J, Marin F, Pergolizzi B, Demarchi B. 2020 Shell palaeoproteomics: First application of peptide mass fingerprinting for the rapid identification of mollusc shells in archaeology. J. Proteomics 227, 103920. (doi:10.1016/j.jprot.2020.103920)

59. Lim MY, Paulo JA, Gygi SP. 2019 Evaluating False Transfer Rates from the Match-between-Runs Algorithm with a Two-Proteome Model. J. Proteome Res. 18, 4020–4026. (doi:10.1021/acs.jproteome.9b00492)

60. Conway JR, Lex A, Gehlenborg N. 2017 UpSetR: an R package for the visualization of intersecting sets and their properties. Bioinformatics 33, 2938–2940. (doi:10.1093/bioinformatics/btx364)

61. Lex A, Gehlenborg N, Strobelt H, Vuillemot R, Pfister H. 2014 UpSet: Visualization of Intersecting Sets. IEEE Trans. Vis. Comput. Graph. 20, 1983–1992. (doi:10.1109/TVCG.2014.2346248)

62. Sharpe JR, Sammons RL, Marquis PM. 1997 Effect of pH on protein adsorption to hydroxyapatite and tricalcium phosphate ceramics. Biomaterials 18, 471–476. (doi:10.1016/S0142-9612(96)00157-3)

63. Stevens SM Jr, Wolverton S, Venables B, Barker A, Seeley KW, Adhikari P. 2010 Evaluation of microwave-assisted enzymatic digestion and tandem mass spectrometry for the identification of protein residues from an inorganic solid matrix: implications in archaeological research. Anal. Bioanal. Chem. 396, 1491–1499. (doi:10.1007/s00216-009-3341-4)

64. Collins MJ, Bishop AN, Farrimond P. 1995 Sorption by mineral surfaces: Rebirth of the classical condensation pathway for kerogen formation? Geochim. Cosmochim. Acta 59, 2387–2391. (doi:10.1016/0016-7037(95)00114-F)

65. Jung F, Frey K, Zimmer D, Mühlhaus T. 2023 DeepSTABp: A Deep Learning Approach for the Prediction of Thermal Protein Stability. Int. J. Mol. Sci. 24. (doi:10.3390/ijms24087444)

66. Charoenkwan P, Ahmed S, Nantasenamat C, Quinn JMW, Moni MA, Lio’ P, Shoombuatong W. 2022 AMYPred-FRL is a novel approach for accurate prediction of amyloid proteins by using feature representation learning. Sci. Rep. 12, 1–14. (doi:10.1038/s41598-022-11897-z)

67. Erdős G, Dosztányi Z. 2020 Analyzing Protein Disorder with IUPred2A. Curr. Protoc. Bioinformatics 70, e99. (doi:10.1002/cpbi.99)

68. Kozlowski LP. 2016 IPC - Isoelectric Point Calculator. Biol. Direct 11, 55. (doi:10.1186/s13062-016-0159-9)

69. Wang S, Li W, Liu S, Xu J. 2016 RaptorX-Property: a web server for protein structure property prediction. Nucleic Acids Res. 44, W430–W435. (doi:10.1093/nar/gkw306)

70. Wang S, Peng J, Ma J, Xu J. 2016 Protein Secondary Structure Prediction Using Deep Convolutional Neural Fields. Sci. Rep. 6, 1–11. (doi:10.1038/srep18962)

71. Buric F, Viknander S, Fu X, Lemke O, Zrimec J, Szyrwiel L, Mueleder M, Ralser M, Zelezniak A. 2023 The amino acid sequence determines protein abundance through its conformational stability and reduced synthesis cost. bioRxiv. [preprint], 2023.10.02.560091. (doi:10.1101/2023.10.02.560091)

72. Pollastri G, Przybylski D, Rost B, Baldi P. 2002 Improving the prediction of protein secondary structure in three and eight classes using recurrent neural networks and profiles. Proteins 47, 228–235. (doi:10.1002/prot.10082)

73. Demarchi B et al. 2016 Protein sequences bound to mineral surfaces persist into deep time. Elife 5, e17092. (doi:10.7554/eLife.17092)

74. Scott A. 2022 Investigation of ancient proteins in archaeological material. PhD, Friedrich Schiller University Jena.

75. Zhang Y, Che H, Li C, Jin T. 2023 Food Allergens of Plant Origin. Foods 12, 2232. (doi:10.3390/foods12112232)

76. Scott A et al. 2021 Exotic foods reveal contact between South Asia and the Near East during the second millennium BCE. Proc. Natl. Acad. Sci. U. S. A. 118, e2014956117. (doi:10.1073/pnas.2014956117)

77. Hilscher A, Heister K, Siewert C, Knicker H. 2009 Mineralisation and structural changes during the initial phase of microbial degradation of pyrogenic plant residues in soil. Org. Geochem. 40, 332–342. (doi:10.1016/j.orggeochem.2008.12.004)

78. Knicker H, Río JC del, Hatcher PG, Minard RD. 2001 Identification of protein remnants in insoluble geopolymers using TMAH thermochemolysis/GC–MS. Org. Geochem. 32, 397–409. (doi:10.1016/S0146-6380(00)00186-8)

79. Knicker H. 2004 Stabilization of N-compounds in soil and organic-matter-rich sediments—what is the difference? Mar. Chem. 92, 167–195. (doi:10.1016/j.marchem.2004.06.025)

80. Knicker H, Hatcher PG. 1997 Survival of Protein in an Organic-Rich Sediment: Possible Protection by Encapsulation in Organic Matter. Naturwissenschaften 84, 231–234. (doi:10.1007/s001140050384)

81. Panettieri M, Knicker H, Murillo JM, Madejón E, Hatcher PG. 2014 Soil organic matter degradation in an agricultural chronosequence under different tillage NMR. Soil Biol. Biochem. 78, 170–181. (doi:10.1016/j.soilbio.2014.07.021)

82. del Río JC, Olivella MA, Knicker H, de las Heras FXC. 2004 Preservation of peptide moieties in three Spanish sulfur-rich Tertiary kerogens. Org. Geochem. 35, 993–999. (doi:10.1016/j.orggeochem.2004.06.002)

83. Ramsøe A, van Heekeren V, Ponce P, Fischer R, Barnes I, Speller C, Collins MJ. 2020 DeamiDATE 1.0: Site-specific deamidation as a tool to assess authenticity of members of ancient proteomes. J. Archaeol. Sci. 115, 105080. (doi:10.1016/j.jas.2020.105080)

84. Tanriver G, Monard G, Catak S. 2022 Impact of Deamidation on the Structure and Function of Antiapoptotic Bcl-xL. J. Chem. Inf. Model. 62, 102–115. (doi:10.1021/acs.jcim.1c00808)

85. Griffin RC, Penkman KEH, Moody H, Collins MJ. 2010 The impact of random natural variability on aspartic acid racemization ratios in enamel from different types of human teeth. Forensic Sci. Int. 200, 148–152. (doi:10.1016/j.forsciint.2010.04.005)

86. Cao B, Porollo A, Adamczak R, Jarrell M, Meller J. 2005 Enhanced recognition of protein transmembrane domains with prediction-based structural profiles. Bioinformatics 22, 303–309. (doi:10.1093/bioinformatics/bti784)

87. Barnard H et al. 2007 Mixed results of seven methods for organic residue analysis applied to one vessel with the residue of a known foodstuff. J. Archaeol. Sci. 34, 28–37. (doi:10.1016/j.jas.2006.03.010)

88. Heaton K, Solazzo C, Collins MJ, Thomas-Oates J, Bergström ET. 2009 Towards the application of desorption electrospray ionisation mass spectrometry (DESI-MS) to the analysis of ancient proteins from artefacts. J. Archaeol. Sci. 36, 2145–2154. (doi:10.1016/j.jas.2009.05.016)

89. Perez-Riverol Y et al. 2022 The PRIDE database resources in 2022: a hub for mass spectrometry-based proteomics evidences. Nucleic Acids Res. 50, D543– D552. (doi:10.1093/nar/gkab1038)

