## supplementary text for "The impact of cooking and burial on proteins: a characterisation of experimental foodcrusts and ceramics"

#### S1: Materials and method

**Sample creation**

Experiments were undertaken outdoors at the YEAR Centre at the University of York, UK. The experiment was designed to simulate cooking over an open fire. A portion of each food type (red deer meat, Atlantic salmon flesh and ground chestnuts) was separately placed into a replica pot, covered with water and heated on an open fire. Temperature was measured on the outside of each vessel using a thermocouple. The pots were left to boil for 1 hour, with water regularly refilled. Subsequently, each pot was emptied and reused for another 1 hour in the same manner. This action was repeated five times for the chestnut flour and 15 times for the deer and salmon [Table S2, 1]. This experiment was replicated in triplicate, with three pots for each ingredient. Subsequently, each pot was split in half: one half was directly analysed for APAAs, the other was buried for six months (May–November 2018) at the YEAR Centre (latitude 53.95; longitude –1.09; soil pH 7.16) prior to analysis for APAAs. All animal products used were acquired commercially (i.e. from a butcher and fishmonger).

**Protein extraction**

0.5M EDTA was added to 100mg of each sample which was then rotated for 5 days. Subsequently, samples were centrifuged at 14000 RCF for 10 minutes and 400 µL of supernatant placed in a new microcentrifuge tube to which 200 µL of 6M GuHCl was also added, followed by 30 µL of 100mM CAA/100 mM TCEP. Samples were briefly vortexed, then place in thermomixer at 99◦C for 10 minutes, then allowed to cool. 20 µl of a 20 µg/µl aqueous mixture containing 1:1 hydrophobic:hydrophilic SpeedBeads™ magnetic carboxylate-modified particles (Sigma Aldrich, Massachusetts, USA, Catalog Nos. GE45152105050250 and GE65152105050250) was then added to each sample and which was mixed by gentle resuspension. 350 µL of absolute ethanol was added to each sample and vortexed to mix. Samples were incubated in the thermomixer at 24◦C with 1000 RPM of shaking for 5 minutes. Samples were then placed on the magnetic rack and allowed to fully migrate for at least 2 minutes. With the magnet still in place, liquid was removed to waste without disturbing the beads. The magnet was removed from the rack and 200 µL of 80% ethanol added to each sample, prior to resuspending the beads in the solution. This wash step was repeated 3 times in total. 100 µL of a 0.04 µg/µL sequencing grade modified trypsin (Promega, Wisconsin, USA, Catalog No. V5111) solution was added to each sample, followed by gentle resuspension. Samples were incubated at 37◦C for 18 hours with 750 RPM of mixing. Samples were placed on a magnetic rack, and beads allowed to migrate for at least 2 minutes. The supernatant (extract) was then carefully removed and placed in a new microcentrifuge tube. The digestion was stopped by adding 10 µL of 5% v/v TFA, reducing pH below 2.5. Samples were then desalted using C18 ziptips (Pierce, Massachusetts, USA, Catalogue No. 87784).

**Machine analysis**

Peptides were re-suspended in aqueous 0.1% trifluoroacetic acid (v/v) then loaded onto an mClass nanoflow UPLC system (Waters) equipped with a nanoEaze M/Z Symmetry 100 Å C 18 , 5 µm trap column (180 µm x 20 mm, Waters) and a PepMap, 2 µm, 100 Å, C 18 EasyNano nanocapillary column (75 m x 500 mm, Thermo). The trap wash solvent was aqueous 0.05% (v:v) trifluoroacetic acid and the trapping flow rate was 15 µL/minute. The trap was washed for 5 minutes before switching flow to the capillary column. Separation used gradient elution of two solvents: solvent A, aqueous 0.1% (v:v) formic acid; solvent B, acetonitrile containing 0.1% (v:v) formic acid. The flow rate for the capillary column was 300 nL/minute and the column temperature was 40°C. The linear multi-step gradient profile was: 3-10% B over 7 minutes, 10-35% B over 30 minutes, 35-99% B over 5 minutes and then proceeded to wash with 99% solvent B for 4 minutes. The column was returned to initial conditions and re-equilibrated for 15 minutes before subsequent injections.

The nanoLC system was interfaced with an Orbitrap Fusion Tribrid mass spectrometer (Thermo) with an EasyNano ionisation source (Thermo). Positive ESI-MS and MS 2 spectra were acquired using Xcalibur software (version 4.0, Thermo). Instrument source settings were: ion spray voltage, 1900-2100 V; sweep gas, 0 Arb; ion transfer tube temperature; 275°C. MS 1 spectra were acquired in the Orbitrap with: 120,000 resolution, scan range: m/z 375-1,500; AGC target, 4e 5 ; maximum fill time, 100 ms. Data dependant acquisition was performed in top speed mode using a 1 s cycle, selecting the most intense precursors with charge states >1. Easy-IC was used for internal calibration. Dynamic exclusion was performed for 50 s post precursor selection and a minimum threshold for fragmentation was set at 5e3. MS 2 spectra were acquired in the linear ion trap with: scan rate, turbo; quadrupole isolation, 1.6 m/z; activation type, HCD; activation energy: 32%; AGC target, 5e 3 ; first mass, 110 m/z; maximum fill time, 100 ms. Acquisitions were arranged by Xcalibur to inject ions for all available parallelizable time.

**Data filtering**

The data was filtered to remove potential machine carry-over and laboratory contaminants. All proteins derived from cRAP (the contaminants database) were removed prior to downstream analysis. An exception was made for peptides matching to Uniprot accessions P68082 (Horse Myoglobin) and P00004 (Horse Cytochrome C) which were found in deer samples, and were likely deer peptides which had matched to the similar horse sequence present in cRAP. All peptides found in laboratory negatives were removed from all samples in downstream analysis. Following common laboratory contaminants such as human keratin, peptides present in lab blanks were primarily derived from salmon (although all at low numbers of identified peptides, ≤2). While a small amount of cross-contamination from the field cooking experiments is evident, we do not believe that this detracts from the utility of this experiment given that the sample creation attempted to replicate potential Mesolithic cooking practices, with pots containing different ingredients processed side-by-side on an open fire. Some carry-over occurred during LC-MS/MS analysis in samples following those with high levels of protein. This carry-over was removed using a custom-made python script which is described below.

**Machine carry-over screening**

In this study, given the relatively high numbers of MS injections and the acquisition of samples of both high and low protein yield, we carefully scrutinised instrument carry-over during LC-MS/MS data acquisition. Examining instrument washes run between each sample, carry-over was observed following protein rich samples such as the fresh ingredients and unburied food crusts (Figure S1). When individual peptides were investigated, the carry-over followed a first-order decay loss with a constant rate of approximately k=0.5 (based on the peptides for which there was sufficient data to calculate k), meaning that the decay of a peptide derived from the contaminating sample was expected to be present at a concentration of 60.653% in any subsequent sample, relative to the concentration of that peptide in the proceeding machine wash.

To mitigate the impact of this carry-over on the data, a python script was written to remove all peptides identified in a given machine wash from the following sample, if at a concentration of that peptide was above a specified threshold (https://github.com/miranda-e/deblanker). For a peptide to pass the threshold (ie. thus to be seen as genuinely present in a sample), it had to have a concentration of at least 121.31% compared to the preceding wash This concentration indicated that a peptide was presented in twice the concentration expected from first order decay with a constant of 0.5 and the peptide was was genuinely present in the sample. The proportion of peptides which did not pass the threshold (and were thus removed from further analysis can be seen in Figure S1. Despite this careful treatment, it is possible that very low-protein content samples located immediately in the wake of carry-over events may have the highest potential to give false-positives. The samples with the highest risk have been annotated with an asterisk in Supplementary Table 1 and arrows in Figure S1.

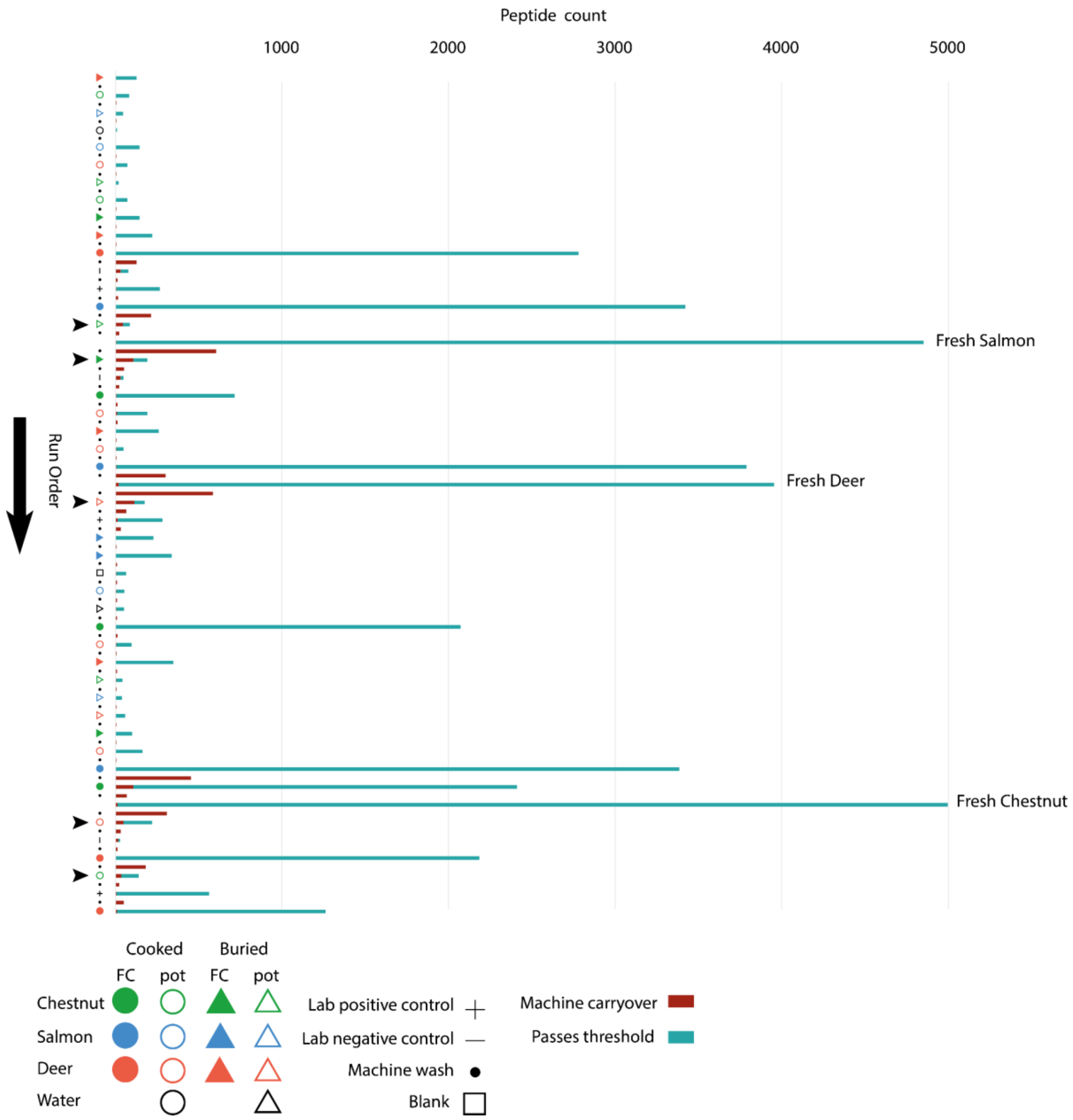


**Figure S1: Peptide abundance (count) in each sample organised in order of machine run (left to right). Blue indicates peptides which passed the threshold and were seen as genuine, red indicates peptides which did not pass the threshold and were not seen as genuinely present in the sample. Arrows indicate samples at highest risk of carry-over.**

### S2: Experimental samples

| Lab sample code | Bondetti et al. 2021 Sample code | Sample type | Cooking stage | Buried/Unburied | Ingredient |
|---|---|---|---|---|---|
| M01 | - | Ingredient | Fresh | Unburied | Deer |
| M02 | - | Ingredient | Fresh | Unburied | Salmon |
| M03 | - | Ingredient | Fresh | Unburied | Chestnut |
| M05 | EDA1P | Ceramic | Cooked | Unburied | Deer |
| M06 | EDA1C | Foodcrust | Cooked | Unburied | Deer |
| M07 | EDB1P | Ceramic | Cooked | Unburied | Deer |
| M08 | EDB1C | Foodcrust | Cooked | Unburied | Deer |
| M09 | EDC1P | Ceramic | Cooked | Unburied | Deer |
| M10 | EDC1C | Foodcrust | Cooked | Unburied | Deer |
| M11 | EFA1P | Ceramic | Cooked | Unburied | Salmon |
| M12 | EFA1C | Foodcrust | Cooked | Unburied | Salmon |
| M13 | EFB1P | Ceramic | Cooked | Unburied | Salmon |
| M14 | EFB1C | Foodcrust | Cooked | Unburied | Salmon |
| M15 | EFC1P | Ceramic | Cooked | Unburied | Salmon |
| M16 | EFC1C | Foodcrust | Cooked | Unburied | Salmon |
| M17 | ECA1P | Ceramic | Cooked | Unburied | Chestnut |
| M18 | ECA1C | Foodcrust | Cooked | Unburied | Chestnut |
| M19 | ECB1P | Ceramic | Cooked | Unburied | Chestnut |
| M20 | ECB1C | Foodcrust | Cooked | Unburied | Chestnut |
| M21 | ECC1P | Ceramic | Cooked | Unburied | Chestnut |
| M22 | ECC1C | Foodcrust | Cooked | Unburied | Chestnut |
| M23 | BLK1 | Ceramic | Boiled | Unburied | Water |
| M24 | EDA2P | Ceramic | Cooked | Buried | Deer |
| M25 | EDA2C | Foodcrust | Cooked | Buried | Deer |
| M26 | EDB2P | Ceramic | Cooked | Buried | Deer |
| M27 | EDB2C | Foodcrust | Cooked | Buried | Deer |
| M28 | EDC2P | Ceramic | Cooked | Buried | Deer |
| M29 | EDC2C | Foodcrust | Cooked | Buried | Deer |
| M30 | EFA2P | Ceramic | Cooked | Buried | Salmon |
| M31 | EFA2C | Foodcrust | Cooked | Buried | Salmon |

| | | | | | |
|---|---|---|---|---|---|
| M32 | EFB2P | Ceramic | Cooked | Buried | Salmon |
| M33 | EFB2C | Foodcrust | Cooked | Buried | Salmon |
| M34 | EFC2P | Ceramic | Cooked | Buried | Salmon |
| M35 | EFC2C | Foodcrust | Cooked | Buried | Salmon |
| M36 | ECA2P | Ceramic | Cooked | Buried | Chestnut |
| M37 | ECA2C | Foodcrust | Cooked | Buried | Chestnut |
| M38 | ECB2P | Ceramic | Cooked | Buried | Chestnut |
| M39 | ECB2C | Foodcrust | Cooked | Buried | Chestnut |
| M40 | ECC2P | Ceramic | Cooked | Buried | Chestnut |
| M41 | ECC2C | Foodcrust | Cooked | Buried | Chestnut |
| M42 | EBLK2P | Ceramic | | Buried | Boiled |
| M04 | BLK0 | Ceramic | Unused | Powdered | |
| M43 | | Mould | Mould | | |

### S3: Protein and peptide count and buried foodcrust lowest common ancestor

**Number of peptides and proteins per buried foodcrust replicate, and most specific LCA for each replicate**

| Ingredient | Chestnut | | Deer | | Salmon | |
|---|---|---|---|---|---|---|
| Replicate | Peptides | Proteins | Peptides | Proteins | Peptides | Proteins |
| 1 | 39 | 25 | 137 | 75 | 203 | 54 |
| 2 | 23 | 16 | 80 | 47 | 193 | 49 |
| 3 | 21 | 14 | 151 | 73 | 157 | 66 |
| | Lowest Common Ancestor | | | | | |
| 1 | Fagaceae/Quercus lobata | | Cervus elaphus hippelaphus | | Salmo | |
| 2 | Fagaceae/Quercus lobata | | Cervidae | | Salmo Salar | |
| 3 | N/A | | Cervidae | | Salmo Salar | |

### S4: Characterisation background and methodology

**Bulk amino acid count**

Amino acid count was investigated as a potential characteristic relating to peptide preservation. Peptide sequence is responsible for the individual characteristics of each protein. Therefore amino acid abundance may be a useful metric to understand, on a basic level, if there are any notable differences between the peptides that survive under the different experiment conditions. Total quantity of each amino acid was calculated per replicate, and compiled on Figure 5a.

**Peptide length**

Peptide length was investigated as a potential indicator of diagenesis from the cooking and entrapment process and/or from burial. Peptide length for all peptides was compiled for each replicate and compiled on Figure 5b.

**Peptide hydropathicity**

We investigated whether peptide hydropathicity (solubility) correlated with peptide abundance in fresh, cooked or buried samples, with the hypothesis that less water soluble (hydrophobic) peptides might preferentially survive in buried samples. Grand Average of Hydropathy (GRAVY) score is a standard measure of protein polarity, which is calculated by summing the hydropathy values for all amino acids in a protein then dividing by the total number of amino acids. A negative GRAVY value indicates that the protein is non-polar and positive value indicates that the protein is polar. GRAVY scores were calculated on a peptide level for each peptide in the fresh, cooked food crust and buried foodcrust samples and their density plotted, to investigate the impact of hydropathicity on peptide preservation (Figure 5c).

#### Peptide isoelectric point

Isoelectric point (pI) is the pH at which a given protein or peptide has an overall charge of zero. A peptide is considered acidic if it has a pI <7, and basic if it has a pI > 7 [2]. An isoelectric point calculator created by Kozlowski [3] was used to calculate the isoelectric point for all identified peptides. These were then plotted by density (peptide count) to investigate the distribution of acidic and basic peptides in fresh ingredients, cooked foodcrust and buried foodcrusts (Figure 5d).

#### Protein thermal stability

Thermal stability is the ability of proteins to resist changes in structure caused by heating. We investigated this characteristic with the hypothesis that thermostable peptides would persist through the cooking, entrapment and burial process. Melting temperature (Tm) is often used as a measure of protein thermal stability. Tm is the temperature at which the concentration of the protein in its folded state equals the concentration of the protein in the unfolded state. DeepSTABp was used to calculate the Tm of all proteins [4]. These were then plotted by density (peptide count) to investigate the distribution of melting temperature for proteins with surviving peptides in all replicates (Figure 5g).

#### Protein amyloid propensity prediction

The propensity for proteins to form amyloid fibres was investigated as a potential aiding factor in protein preservations in buried foodcrust samples, with the hypothesis that amyloids form a fibril-like structure that may be resistant to diagenesis. The amyloid propensity for each protein was generated using AMYPred-FRL [5]. Amyloid probability score generated by AMYPred-FRL for each protein was plotted density (peptide count) for each sample (fresh, cooked foodcrust, buried foodcrust) (Figure 5h).

#### Peptide secondary structure

Secondary structure is innately linked to a protein's function and stability. Proteins form a variety of secondary structures dependent on hydrogen bonds formed between atoms of the polypeptide backbone. Secondary structure was investigated through the application of “Predict_Property”, a standalone, offline version of RaptorX web server [6,7] (https://github.com/realbigws/Predict_Property). Predict property was used to protect the eight-state secondary structure (SS8) secondary structure of all proteins present in the filtered sample using the following categories: G: 310 helix, H: alpha-helix, I: pi-helix, E: beta-strand, B:beta-bridge, T: beta-turn, S: high curvature loop, and L for irregular. A python script was then written to match peptides to the correct position within a sequence, and extract the SS8 DSSR secondary structure at that point. These secondary structures were then analysed for overall abundance within each experiment, to understand whether any particular secondary structure was more likely to preserve under each condition (Figure 5f).

#### Relative solvent accessibility

Relative solvent accessibility (RSA) is a measure of the extent of exposure or burial of an amino acid in a protein in the 3D structure, and therefore how accessible that residue is to a solvent. Both the location in the protein's native state, and the characteristics of each individual amino acid impact RSA. RSA was calculated using Predict Property, which is described above. These RSAs were then analysed for overall abundance within each experiment, to understand whether any particular secondary structure was more likely to preserve under each condition.

#### Disorder prediction

Intrinsic disorder was investigated as an alternative way of investigating the impact of structural flexibility on protein preservation. Intrinsically disordered proteins are those that do not have a well-defined tertiary structure and are thus usually flexible. Disordered proteins hold important roles, often related to their flexibility. IUpred2A was used to calculate the disorder propensity of all proteins present in the samples [8]. A python script was then created which extracted the average disorder propensity of each peptide, which are compiled in Figure 5e.

#### Deamidation

Variable Post Translational Modifications (PTMS) included in the Maxquant Search included oxidation (M), acetylation (protein N-term), deamidation (NQ). Proportion of deamidated peptides compared to non-deamidated peptides was compiled for each experiment (combined replicates) on Figure 6. Unfortunately, due to the necessary peptide filtration steps discussed below, it was not possible to apply deamiDATE here to assess site-specific deamidation. Further investigation of site-specific PTMs may reveal evidence for aspects of food preparation and taphonomy.

### S5: Lipid content

We compared the protein results to lipid quantities generated from FID lipid quantities generated by Bondetti et al.'s [1] analysis of the same samples. This was plotted against total peptide and protein count, the results of which can be seen below.

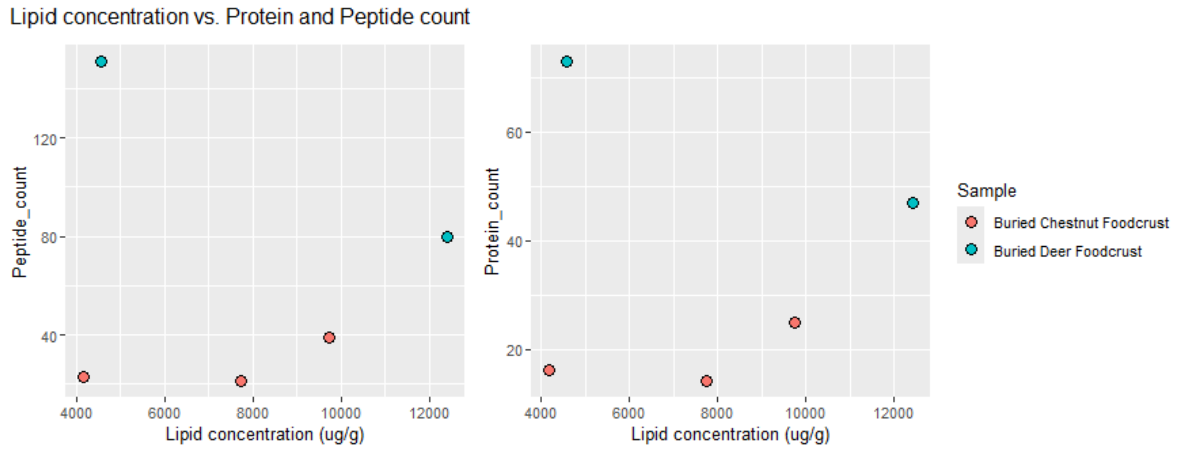


| Sample | Replicate | Peptides | Proteins | Lipid concentration (ug/g) |
|---|---|---|---|---|
| Buried Deer Foodcrust | 2 | 80 | 47 | 12409.4 |
| Buried Deer Foodcrust | 3 | 151 | 73 | 4565.73 |
| Buried Chestnut Foodcrust | 1 | 39 | 25 | 9738.47 |
| Buried Chestnut Foodcrust | 2 | 23 | 16 | 4169 |
| Buried Chestnut Foodcrust | 3 | 21 | 14 | 7733.56 |

### S6: Percentage of proteins annotated with gene ontology terms, by experiment

| Experiment | Percentage of proteins which are annotated with gene ontology terms |
|---|---|
| buried_cooked_chestnut_pot | 53.84615 |
| buried_cooked_deer_foodcrust | 64.02116 |
| buried_cooked_deer_pot | 57.57576 |
| buried_cooked_salmon_foodcrust | 41.77215 |
| buried_cooked_salmon_pot | 84 |
| cooked_chestnut_foodcrust | 39.07929 |
| cooked_chestnut_pot | 80 |
| cooked_deer_foodcrust | 50.23451 |
| cooked_deer_pot | 65.04854 |
| cooked_salmon_foodcrust | 33.67987 |
| cooked_salmon_pot | 77.77778 |
| fresh_chestnut | 42.79079 |
| fresh_deer | 61.70878 |
| fresh_salmon | 35.2383 |

### S7: Field experiment carryover (peptide proportion) by sample

| Ingredient | Matrix | Buried/unburied | Cooking | Replicate | Percentage of peptide count derived from non-target ingredients | Accepted for protein property analysis |
|---|---|---|---|---|---|---|
| Chestnut | Ingredient | Unburied | Fresh | 1 | 0.81 | True |
| Chestnut | Foodcrust | Unburied | Cooked | 1 | 1.83 | True |
| Chestnut | Foodcrust | Unburied | Cooked | 2 | 1.67 | True |
| Chestnut | Foodcrust | Unburied | Cooked | 3 | 1.46 | True |
| Chestnut | Foodcrust | Buried | Cooked | 1 | 10.26 | False |
| Chestnut | Foodcrust | Buried | Cooked | 2 | 34.78 | False |
| Chestnut | Foodcrust | Buried | Cooked | 3 | 28.57 | False |
| Deer | Ingredient | Unburied | Fresh | 1 | 1.15 | True |
| Deer | Foodcrust | Unburied | Cooked | 1 | 1.17 | True |
| Deer | Foodcrust | Unburied | Cooked | 2 | 0.90 | True |
| Deer | Foodcrust | Unburied | Cooked | 3 | 1.05 | True |
| Deer | Foodcrust | Buried | Cooked | 1 | 1.41 | True |
| Deer | Foodcrust | Buried | Cooked | 2 | 0.00 | True |
| Deer | Foodcrust | Buried | Cooked | 3 | 1.30 | True |
| Salmon | Ingredient | Unburied | Fresh | 1 | 0.78 | True |

| | | | | | | |
|---|---|---|---|---|---|---|
| Salmon | Foodcrust | Unburied | Cooked | 1 | 0.36 | True |
| Salmon | Foodcrust | Unburied | Cooked | 2 | 0.34 | True |
| Salmon | Foodcrust | Unburied | Cooked | 3 | 0.30 | True |
| Salmon | Foodcrust | Buried | Cooked | 1 | 0.99 | True |
| Salmon | Foodcrust | Buried | Cooked | 2 | 1.55 | True |
| Salmon | Foodcrust | Buried | Cooked | 3 | 1.91 | True |

#### Bibliography


1. Bondetti M, Scott E, Courel B, Lucquin A, Shoda S, Lundy J, Labra-Odde C, Drieu L, Craig OE. 2021 Investigating the formation and diagnostic value of ω -( o -alkylphenyl)alkanoic acids in ancient pottery. *Archaeometry* **63**, 594–608.

2. Moldoveanu SC, David V. 2016 Chapter 5 - Properties of Analytes and Matrices Determining HPLC Selection In: Serban C. Moldoveanu, Victor David, editors. Selection of the HPLC Method in Chemical Analysis: Elsevier, 2016, 189–230.

3. Kozlowski LP. 2016 IPC - Isoelectric Point Calculator. *Biol. Direct* **11**, 55.

4. Jung F, Frey K, Zimmer D, Mühlhaus T. 2023 DeepSTABp: A Deep Learning Approach for the Prediction of Thermal Protein Stability. *Int. J. Mol. Sci.* **24**, 74444 (doi:10.3390/ijms24087444)

5. Charoenkwan P, Ahmed S, Nantasenamat C, Quinn JMW, Moni MA, Lio' P, Shoombuatong W. 2022 AMYPred-FRL is a novel approach for accurate prediction of amyloid proteins by using feature representation learning. *Sci. Rep.* **12**, 1–14.

6. Wang S, Li W, Liu S, Xu J. 2016 RaptorX-Property: a web server for protein structure property prediction. *Nucleic Acids Res.* **44**, W430–W435.

7. Wang S, Peng J, Ma J, Xu J. 2016 Protein Secondary Structure Prediction Using Deep Convolutional Neural Fields. *Sci. Rep.* **6**, 1–11.

8. Erdős G, Dosztányi Z. 2020 Analyzing Protein Disorder with IUPred2A. *Curr. Protoc. Bioinformatics* **70**, e99.
